# Phenotypic and genomic changes associated with the evolutionary loss of heterostyly in primroses

**DOI:** 10.64898/2026.10.19.760172

**Authors:** Barbara Keller, Elena Conti, Hassan Mansour, Emiliano Mora-Carrera, Rebecca L. Stubbs, Narjes Yousefi, Giacomo Potente

## Abstract

- Trait loss has been documented across broad phylogenetic scales and dissected genetically within species, but how the genetic basis of loss varies among lineages and over evolutionary time remains poorly understood. Heterostyly, a floral polymorphism, represents an ideal case study, as it has been lost repeatedly across angiosperms. In *Primula*, heterostyly is controlled by the *S*-locus supergene, and loss-of-function mutations in *CYP*ᵀ cause transitions to homostyly.
- We investigated the critically endangered *Primula boveana*, whose floral morphology is debated, in the diploid, self-compatible *P.* sect. *Sphondylia*. We quantified herkogamy dynamics during anthesis, performed controlled pollination experiments, assembled a chromosome-scale genome, compared *S*-locus structure and gene expression with heterostylous relatives, and reconstructed phylogenetic relationships within the section.
- Our results revealed that *P. boveana* is homostylous, with a disrupted *CYPᵀ*. Herkogamy declines during anthesis, resulting in early anther-stigma contact, autonomous selfing, and reproductive assurance. Thus, unlike other homostylous *Primula* species, these developmental changes do not represent a delayed selfing strategy.
- These findings link developmental changes in herkogamy with autonomous selfing and support *CYPᵀ* inactivation as the genetic basis of long homostyly in *P. boveana*. The extensive degeneration of *CYPᵀ* suggests that prolonged relaxed selection promotes progressive degeneration of non-functional *S*-genes following heterostyly loss.

## Introduction

Phenotypic traits can be gained or lost as selective pressures change. Traits once advantageous may be lost when they become costly or maladaptive (Porter & Crandall, 2003; Lahti *et al*., 2009). Because many independent loss-of-function mutations can disrupt a trait, whereas trait gain typically requires specific gain-of-function changes, trait loss occurs more frequently than trait gain (Rosenblum *et al*., 2010; Ellers *et al*., 2012; Jones *et al*., 2020; Mora-Carrera *et al*., 2023; 2024; Stone & Wessinger, 2024). An illustrative case in angiosperms comes from heterostyly, a discrete floral heteromorphism found across 28 angiosperm families (Ganders, 1979; Naiki, 2012; Barrett, 2019), which has been lost multiple times independently (Simón-Porcar *et al*., 2024).

Heterostylous species comprise populations with two or three plant types, distinguished by flowers with a reciprocal arrangement of female (stigma) and male (anthers) reproductive organs (Barrett, 2002). Heterostyly promotes outcrossing while minimizing selfing and sexual interference (Barrett, 2002; Keller *et al*., 2014). The loss of heterostyly results in homostyly, in which populations are monomorphic and flowers bear anthers and stigmas in close spatial proximity. This transition typically co-occurs with the loss of self-incompatibility, or, rarely, proceeds *via* purely morphological shifts in already self-compatible lineages (Raimondeau *et al*., 2024; Zeng *et al*., 2024), ultimately enabling seed production without pollinators or mates (i.e., autonomous selfing) (Piper & Charlesworth, 1986; De Vos *et al*., 2012; Yuan *et al*., 2023; Zeng *et al*., 2024).

Autonomous selfing in homostyles may facilitate colonization and persistence in isolation, or reproduction within harsh environments where depauperate pollinator fauna, mate scarcity, and poor habitat suitability restrict outcrossing opportunities (Baker, 1967; Pannell *et al*., 2015). However, selfing also reduces genetic variation in populations, limiting their capacity to adapt to environmental changes (Charlesworth, 2003). Moreover, selfed offspring are often less fit than outcrossed offspring (*i.e*., inbreeding depression), as selfing increases homozygosity at loci with both recessive, deleterious mutations and heterozygote advantage (Baker, 1966; Charlesworth & Charlesworth, 1987). These fitness costs can be mitigated when autonomous selfing occurs only after opportunities for outcrossing have been exhausted, a strategy termed delayed selfing (Lloyd, 1992). For example, some homostylous species such as *Primula halleri* exhibit pronounced anther-stigma separation (i.e., herkogamy) when flowers open, favoring cross-pollination, but this separation diminishes as flowers age, enabling delayed autonomous selfing, an advantage under fluctuating environmental conditions (De Vos *et al*., 2012). However, *P. halleri* is the only homostylous species ever described to possess such a developmental variation in herkogamy, thus it remains unknown whether it is widespread across homostylous species and, if so, whether it is always accompanied by delayed selfing.

The classic model for heterostyly is *Primula* L (primroses; Darwin, 1877), comprising dimorphic populations of typically self-incompatible plants that bear either long styled flowers with low anthers (L-morph, or pins) or short styled flowers with high anthers (S-morph, or thrums; **Fig. S1a**). Ancillary features, such as pollen-size dimorphism, are associated to the floral morphs (Ganders, 1979; Dulberger, 1992). Heterostyly is controlled by a supergene (i.e., a cluster of tightly linked genes controlling complex phenotypes inherited as a single unit due to suppressed recombination; Schwander *et al*., 2014; Thompson & Jiggins, 2014; Berdan *et al*., 2022) known as the *S*-locus, which is hemizygous in thrums (*S/s*) and absent from pins (*s/s*) (Huu *et al*., 2016; Li *et al*., 2016; Potente *et al*., 2022a). The primrose *S*-locus comprises four “core” genes (*CYP^T^*, *GLO^T^*, *PUM^T^*, and *KFB^T^*; “*S*-genes” hereafter), present in the dominant *S*-haplotype but absent from the recessive *s*-haplotype (Potente *et al*., 2022a; 2024). Of them, *GLO^T^* determines high anthers, whereas *CYP^T^* determines short styles and, in self-incompatible species, also female self-incompatibility (Huu *et al*., 2016; 2020; 2022).

Heterostyly is ancestral in *Primula* and was lost multiple times independently both within and between species (Mast *et al*., 2006; De Vos *et al*., 2014), resulting in homostylous, self-compatible flowers (**Fig. S1b**). In all cases investigated, the loss of heterostyly is associated with loss-of-function mutations in *CYP^T^* (Huu *et al*., 2016; 2022). In the heterostylous *P. vulgaris*, homostylous individuals evolved repeatedly *via* nonsynonymous mutations and/or small indels disrupting *CYP^T^*(Mora-Carrera *et al*., 2023; 2024). In the long-homostylous *Primula grandis*, *CYP^T^* shows a more extensive accumulation of mutations (Mora-Carrera *et al*., 2025). In *Hottonia*, the sister clade of *Primula*, the homostylous *H. inflata* has lost *CYP*ᵀ entirely (Potente *et al*., 2026). This progression in *CYP*ᵀ degeneration – from few mutations within species, to extensive mutations or gene loss between species – is consistent with the prediction that relaxed selection accumulates mutations over time (Collin & Miglietta, 2008; Lahti *et al*., 2009; Cheetham *et al*., 2020). As in *Primula*, the transition to homostyly in other angiosperms has repeatedly involved disruption of the gene controlling style length [*Fagopyrum* (Fawcett *et al*., 2023), *Turnera* (Matzke *et al*., 2021), and *Chrysojasminum* (Raimondeau *et al*., 2024)], with the only exception of *Linum trigynum* (Linaceae), where heterostyly loss is associated with downregulation of the gene controlling style length rather than loss-of-function mutations (Gutiérrez-Valencia et al., 2024). Collectively, these cases show that heterostyly loss can arise through diverse molecular changes, from gene downregulation, to loss-of-function mutations, to gene loss.

In *Primula*, homostyly frequently co-occurs with polyploidy (Naiki, 2012), as in *P. grandis* (Mora-Carrera *et al*., 2025), complicating the distinction between genomic changes linked to trait loss and those associated with whole-genome duplication. A diploid clade comprising both heterostylous and homostylous species would instead allow such changes to be identified directly. One such group is *P.* sect. *Sphondylia* (Duby) Rupr., a small (eight species), relatively young diploid clade (ca. 5 Ma; De Vos *et al*., 2014; 2n = 18) that includes both heterostylous and homostylous species. Interestingly, all species in sect. *Sphondylia* are self-compatible (Al Wadi & Richards, 1993; Richards, 2003; De Vos *et al*., 2014), unlike most other heterostylous primroses, which are typically self-incompatible (Richards, 2003). This makes the section suitable for isolating the genetic basis of homostyly both from the confounding effects of polyploidy. Within *P.* sect. *Sphondylia*, a chromosome-scale assembly is already available for the heterostylous *P. edelbergii* (Potente *et al*., 2024), providing a reference for *S*-locus comparison within the section. In *P. edelbergii*, the 634-kb *S*-locus contains the four core *S*-genes, including *CYP^T^*, which was expressed in floral tissues (Potente *et al*., 2024). Notably, since *CYP^T^* controls both style length and female self-incompatibility in thrums of other heterostylous, self-incompatible primroses, its intact function in *P. edelbergii* is consistent with the heterostylous phenotype but seemingly at odds with the self-compatibility of this species (Wedderburn & Richards, 1990; 1992; Al Wadi & Richards, 1993), a paradox that may implicate alterations in the as-yet-uncharacterized male side of the incompatibility system. Together, the diploidy, self-compatibility, and genomic resources of *P.* sect. *Sphondylia* make it an ideal system for identifying the *S*-locus changes underlying heterostyly loss independently of polyploidy and self-incompatibility.

A intriguing species within *P.* sect. *Sphondylia* is *P*. *boveana* Decne ex Duby, a critically endangered endemic of the Sinai Peninsula, Egypt (**Fig. 1a**; Radford *et al*., 2011; Omar, 2014b), which is highly inbred, likely due to recurrent selfing caused by pollinator scarcity and small population sizes (Jiménez *et al*., 2013). Interestingly, the floral morphology of *P. boveana* has been subject to conflicting interpretations: (Ernst, 1949; Wendelbo, 1961) described it as homostylous, while (Al Wadi & Richards, 1993) as heterostylous with dimorphic style length and pollen size but anthers at the same height in both morphs. Reports of flowers with both protruding and sunken stigmas within the same individual (**Fig. 1b**; Zaghloul *et al*., 2016) argue against true heterostyly, instead suggesting homostyly with variation in herkogamy. Whether *P. boveana* is homostylous or heterostylous remains unknown.

**Figure 1.**
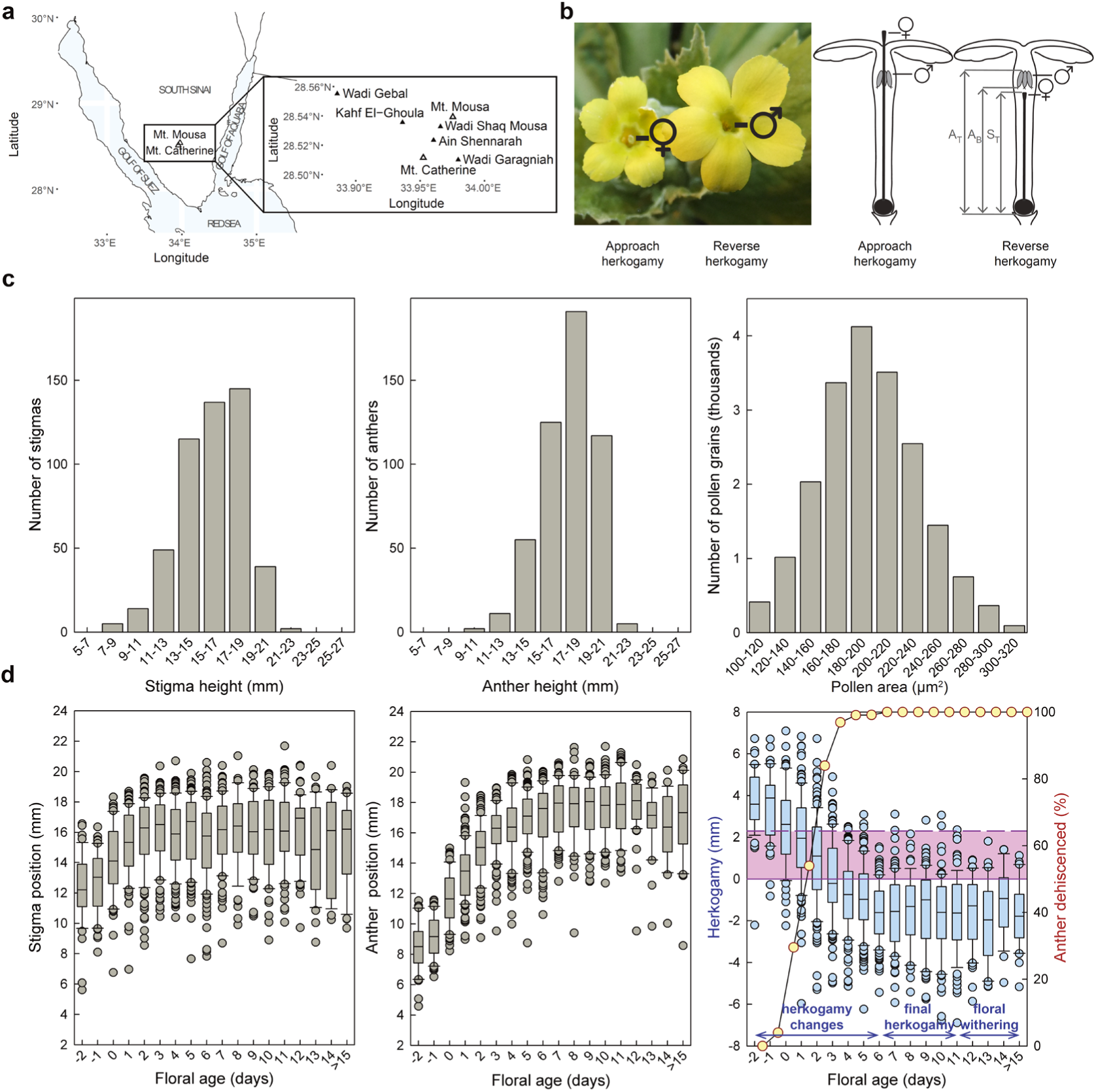
Distribution and variation of floral traits in the endangered *Primula boveana.* (a) *P. boveana* is endemic to the southern Sinai Peninsula of Egypt. Black triangles refer to the five populations known in 2011, located near Wadi Gebal (WG), Khaf El-Ghoula (GO), Wadi Shaq Mousa (SM), Ain Shennerah (AS), and Wadi Garagniah (GR) (see also Jiménez *et al*., 2013). (b) *P. boveana* has flowers with protruding and sunken stigmas on the same plant. Picture **(left panel)** and schematic drawings **(right panel)** of flower with protruding stigma (*i.e.,* approach herkogamy: left) and flower with sunken stigma (*i.e*., reverse herkogamy: right). To assess whether stigmas are positioned above, between, or below anthers, floral traits were measured from the base of the gynoecium to the stigma tip (S_T_), anther base (A_B_), and anther tip (A_T_). (c) *P. boveana* is a homostylous species exhibiting unimodal frequency distributions of anther height (A_T_), stigma height (S_T_), and pollen size. Distributions of anther and stigma height are shown for mature flowers only and reflect final, stable herkogamy after floral development has ceased. Specifically, distributions of (i) anther (A_B_) and (ii) stigma (S_T_) heights for 506 mature flowers (defined as flowers where the positions of anthers and stigmas no longer changed, a condition reached on day 6 after flower opening; see below) and (iii) pollen sizes of 19800 pollen grains. **(d) Variation of floral traits during anthesis.** Boxplots of stigma position (S_T_; **left panel)**, anther position (A_B_; **middle panel)**, herkogamy (S_T_-A_B_; **right panel**; blue), and percentages of flowers with dehisced anthers (red line with yellow circles) across 1387 flowers (for details, see Fig. S2). Floral age on the x axis. Boxplots show the median (horizontal line), interquartile range (box edges, 25th to 75th percentiles), and whiskers extending to the most extreme data points within 1.5 times of the interquartile range (IQR). Data points beyond the whiskers are plotted individually as outliers. *Herkogamy changes* from 2 days before floral opening to 6 days after floral opening, starting with stigma above anthers (approach herkogamy) in most flowers and ending with stigma below anthers (reversed herkogamy) when herkogamy stabilizes on day 6. *Final herkogamy*, with stigma below anthers in most flowers, remains stable from day 6 until the end of anthesis. *Floral withering* starts after day 11 of anthesis. The horizontal purple lines at 0 mm and 2.3 mm indicate the mean anther length (± 0.4 mm standard deviation; data not shown); the pink shaded area between the two horizontal lines thus indicates that stigmas are positioned between the anthers (hence no herkogamy), allowing for autonomous selfing.

Here, we resolved the floral morphology of *P. boveana* and investigated the developmental and genetic basis of its transition to homostyly. First, we tested whether stigma height, anther height, and pollen size are unimodally or bimodally distributed, to determine whether *P. boveana* is homostylous or heterostylous. Having established homostyly with developmental variation in herkogamy, we used controlled pollination experiments to test whether this variation reflects delayed-selfing, the delayed-selfing strategy observed in *P. halleri* (De Vos et al., 2012), or instead results in early, exclusive self-fertilization. Next, we generated a chromosome-scale genome assembly of *P. boveana* and tested whether homostyly in this species is associated with a disruption in *CYP^T^*, as observed in other homostylous Primulaceae (Mora-Carrera *et al*., 2025; Potente *et al*., 2026), despite the self-compatible, diploid background of sect. *Sphondylia*. Finally, to place the molecular changes detected at the *S*-locus in a temporal framework, we resolved the phylogenetic placement of *P. boveana* within sect. *Sphondylia*, allowing us to test whether its loss of heterostyly shares a common origin with that of closely related homostylous species.

Resolving *P. boveana* mating system is important not only for understanding the evolutionary and ecological implications of the transition to homostyly but also, given its critically endangered status, to inform potential future conservation efforts targeting its reproductive biology. More broadly, this study provides a multifaceted account of heterostyly loss, integrating phenotypic, ecological, genomic, and phylogenetic evidence on a transition that has occurred repeatedly across the angiosperms.

## Material and Methods

### A. Study species and plant material

*Primula boveana* (2n = 18) is a critically endangered, rosette-forming perennial restricted to a few small populations in montane wadis above 1700 m a.s.l. in the St. Catherine Protectorate, southern Sinai, Egypt (Moustafa *et al*., 2001; Radford *et al*., 2011; Omar & Elgamal, 2021). Plants grow in moist rock crevices near snow-fed springs, where water is available nearly year-round (Zaghloul *et al*., 2016; Abdelaal *et al*., 2020). Flowers have showy, golden-yellow, hypocrateriform corollas with five lobes and a slender tube bearing five fused anther filaments. Anthers are attached high in the corolla tube. Fruit capsules produce an average of 423 seeds, which accumulate in the soil seed bank and enable survival during dry periods (Moustafa *et al*., 2001; Zaghloul, 2008).

Seeds were collected in 2011 from 35 plants and 11 soil samples across all the five populations known (Moustafa *et al*., 2001; **Fig. 1a, Table S1**);). Seeds were stored at 4°C to break dormancy and germinated in spring 2012 in growth chambers (Sanyo MLR 351H; Panasonic Corp., Japan; 18°C, 55% humidity, 12 h dark at 10°C and 12 h light [22,000 LUX]) at the University of Zurich (UZH), Switzerland. Seedlings were transplanted into individual pots, yielding 289 plants. Of these, 126 and 33 were used in 2012 and 2014, respectively (**Fig. S2a; Table S1**). Plants were cultivated and experiments conducted under standard conditions in a pollinator-free greenhouse at the Botanical Gardens of UZH (18–22°C, ∼65% humidity, 12 h photoperiod, ≥30,000 LUX). Experimental plants were identified by Barbara Keller, and a voucher specimen is deposited at Herbarium Z (Zurich, Switzerland; Z-000195754).

### B. Phenotypic analyses

Floral traits during anthesis (spanning from the day of floral opening, designated as day 0 hereafter, until floral withering), self-pollen deposition, and seed set (after autonomous and manual pollinations) were assessed in partially overlapping subsets of 1826 flowers from 148 plants (2012: 1659 flowers from 126 plants; 2014: 167 flowers from 33 plants; **Fig. S2a**). These data were used to address the following questions: (i) Is *P. boveana* heterostylous or homostylous? (ii) Does herkogamy change during anthesis? (iii) Does developmental variation in herkogamy affect the capacity for autonomous selfing? and (iv) Can autonomous selfing confer reproductive assurance in *P. boveana*?

#### Quantification of floral traits and reproductive traits

Herkogamy was visually assessed in 1,644 intact flowers (126 plants) that appeared ready to open (i.e., initial herkogamy; **Fig. S2a**). Each intact flower was assigned to one of three herkogamy classes: approach initial herkogamy (stigma above anthers), no initial herkogamy (stigma between anthers), and reverse initial herkogamy (stigma below anthers). Subsequently, a subset of 1,387 flowers (125 plants) was randomly harvested (either during the initial assessment period or daily throughout the flowering period until withering) to track floral and reproductive changes (see **Fig. S2b** for daily sample sizes). Flowers were longitudinally dissected, photographed, and preserved in 70% ethanol. The distances from the gynoecium base to stigma tip (S_T_), anther base (A_B_), and anther tip (A_T_) were measured using ImageJ v1.53t (https://imagej.net/ij/; **Fig. 1b**). Herkogamy was calculated as S_T_ - A_B_ (stigma tip minus anther base) and flowers assigned to herkogamy classes (approach: S_T_ > A_T_; no: A_T_ > S_T_ > A_B_; reverse: S_T_ < A_B_). In 92 flowers, anthers were discolored, remained closed, and showed no evidence of pollen release; these flowers were excluded from subsequent analyses requiring pollen release. In the remaining 1,295 flowers, pollen presence on stigmas and anther dehiscence (stomium split and pollen exposed) were assessed and the daily proportion of flowers with dehisced anthers calculated.

Pollen area was determined using photographs of pollen grains mounted in fuchsin jelly (Dafni *et al*., 2005) from one anther each of 68 flowers (33 plants; **Fig. S2a**). A total of 19,800 grains (600 per anther) were automatically quantified using particle analysis in Fiji/ImageJ v1.54f (Schindelin *et al*., 2012). Stigma receptivity was tested in 15 flowers from day 0 to day 6 (**Fig. S2a**) by assessing pollen germination and pollen tube growth after 24–48 h after hand pollination, using Martin’s method (1959) and UV illumination (see **Methods S1**). A stigma was considered receptive if pollen germinated.

To quantify seed set, 257 flowers (70 plants; **Fig. S2a**) were either left intact (126 flowers, 68 plants) to quantify autonomous seed formation or emasculated (131 flowers, 69 plants) to control for unintended pollen transfer. Flowers were bagged after withering, and fruits were harvested at maturity. Seeds and ovules were counted as described in **Methods S1**. Seed set was calculated as the proportion of ovules developing into seeds, based on counts of seeds and total ovules obtained from digital images (**Methods S1**). Ovule number varied substantially among flowers (279–1161 per flower), a pattern previously reported in another homostylous *Primula* (De Vos *et al*., 2012).

We then compared autonomous seed set with seed set from manually-selfed and manually-outcrossed flowers in a pollination experiment (**Fig. S2a**). A total of 167 flowers (33 plants) were randomly assigned to one of four treatments: (i) SELF_AUTO_: intact flowers allowing autonomous self-pollination (41 flowers); (ii) SELF_MANUAL_: emasculated flowers hand-pollinated with pollen from two flowers of the same plant (64); (iii) CROSS: emasculated flowers manually-pollinated with pollen from two flowers of different plants within the same population (53); and (iv) CONTROL: emasculated, unpollinated flowers (9). No CONTROL flowers set seeds, indicating accidental pollination due to flower handling should be rare. Fruits were collected; seeds and ovules were counted as above.

### Analysis of floral traits and reproductive variation

#### (i) Is Primula boveana homostylous?

To determine whether *P. boveana* is heterostylous or homostylous, we analyzed frequency distributions of: (i) stigma and anther heights from 506 mature, non-withered flowers at the stage we termed final herkogamy (i.e., when floral organ positions had stabilized, approximately 6 days after floral opening; **Fig. S2b**); (ii) areas of 19,800 pollen grains (68 flowers; **Fig. S2a**). Distributions were visually assessed for unimodality (consistent with homostyly) or bimodality (indicative of heterostyly). Hartigan’s dip test (*diptest* R package v.0.77.1; Maechler, 2024) was used to test for significant deviation from unimodality.

#### (ii) Does herkogamy change during anthesis?

To evaluate changes in herkogamy during anthesis, we generated daily boxplots of anther, stigma, and herkogamy measurements from 1387 flowers (125 plants) spanning days −2 to +20 of anthesis (**Fig. S2b**). Herkogamy changed from −2 to +6 days and remained stable thereafter (final herkogamy). To test whether anthers and stigmas elevated at different rates, we modeled organ height using a generalized linear mixed-effects model (GLMM) with *organ type* (anther or stigma), *day* (-2 to +6), and their interaction as fixed effects (**Table S2a**). Organ height was square-transformed to improve normality. Model performance was evaluated by inspecting Pearson residuals. Among-plant variation was examined using Spearman rank correlations between *herkogamy* and *day* in 101 plants with ≥ 3 flowers (798 flowers; *ggpubr* R package v.0.6.0; Kassambara, 2023).

#### (iii) Does the capacity for autonomous selfing correlate with initial herkogamy?

To examine the relationship between initial herkogamy and autonomous selfing, we analyzed pollen presence/absence data from 465 flowers (106 plants; **Fig. S2a**). These were selected from 655 mature flowers in which organ positions had stabilized (*i.e*., from day 6 to day 20 [+6 to +20] of anthesis; **Fig. S2b**), excluding 57 flowers with unopened anthers (self-pollen not available), 108 flowers with stigmas positioned between anthers (to avoid contamination), and 25 flowers lacking initial herkogamy data. Of the selected 465 flowers, 134 showed approach, 129 no, and 202 reverse initial herkogamy. Differences among the three classes were assessed using a chi-square test of independence.

To test whether autonomous seed set (proportion of ovules developing into seeds) differed among initial herkogamy classes, we analyzed seed-set data from 126 flowers (68 plants; **Fig. S2a**): 47 showed approach, 54 no, and 25 reverse initial herkogamy. Differences among herkogamy classes were assessed using a GLMM (**Table S2b**). Model performance was evaluated by comparing observed and predicted seed presence/absence and inspecting Pearson residuals against predicted probabilities.

#### (iv) Does autonomous selfing reduce seed set compared to outcrossing?

To assess whether autonomous selfing produces fewer seeds than outcrossing and/or manual selfing, we compared seed-set data from 41 autonomous self-, 64 manual self-, and 53 cross-pollinated flowers using a GLMM (33 plants; **Fig. S2a**; **Table S2c**). Fixed effects included *pollination treatment* (SELF_AUTO_, SELF_MANUAL_, CROSS), *population* (GR, SM, WG), and their interaction. *Population* was included because reproductive output from selfing and outcrossing can vary among populations (Thoen *et al*., 2024). Populations AS and GO were excluded due to insufficient sample sizes. Model performance was evaluated as above.

#### (v) How much reproductive assurance is provided by autonomous selfing?

For each herkogamy class, reproductive assurance was calculated as the product of its frequency at flower opening and the proportion of seed set achieved through autonomous selfing relative to manual self-pollination. Summing these values across herkogamy classes yielded an estimate of reproductive assurance, defined as the proportion of potential seed production maintained in the absence of pollinators.

Statistical analyses were conducted using SPSS v29.0.0.0 (IBM Corp., Armonk, NY, USA), unless otherwise specified. Sequential Bonferroni corrections were applied for multiple comparisons when applicable. All GLMMs accounted for hierarchical data *via* random effects and unbalanced data using Satterthwaite’s method for denominator degrees of freedom.

### C. Genomic analyses

#### Nucleic acid extraction and sequencing

To assemble the genome of *P. boveana*, high molecular weight DNA was extracted from fresh leaves of a single individual (hereafter “reference individual”) from the Wadi Garagniah population using a modified cetyltrimethylammonium bromide (CTAB) protocol (Healey *et al*., 2014; Potente *et al*., 2022a). Long-read sequencing was performed on Oxford Nanopore MinION R9.4.1 flow cells (∼67× coverage), and reads were basecalled with Guppy v6.6.0.1 using the Super High Accuracy (SUP) model. In addition, Illumina paired-end 150-bp whole-genome sequencing (WGS; ∼113× coverage) and Hi-C sequencing for scaffolding were performed at the Functional Genomics Center Zurich (FGCZ, Switzerland) and Arima Genomics (USA), respectively.

RNA was extracted from leaves, flowers, and young and mature floral buds of the reference individual and sequenced on an Illumina NovaSeq 6000 at FGCZ (∼100 million paired-end 150-bp reads per tissue). To validate genomic patterns at the *S*-locus, a second individual from the Wadi Shaq Mousa population was sequenced using Illumina whole-genome resequencing (∼20× coverage). Details are provided in **Methods S2**.

#### Genome assembly and annotation of Primula boveana and synteny with P. edelbergii

Genome size was estimated from k-mer frequencies of the forward Illumina reads using GenomeScope (http://qb.cshl.edu/genomescope/; Vurture *et al*., 2017). Illumina and Nanopore reads were combined for *de novo* assembly with MaSuRCA v4.0.5 (Zimin *et al*., 2013), with Flye v2.5 (Kolmogorov *et al*., 2019) used for final contigging. Contigs were polished with POLCA (Zimin & Salzberg, 2020) and scaffolded with Hi-C data using YaHS v1.2a.1 (Zhou *et al*., 2022). Assembly quality was assessed with KAT (Kmer Analysis Toolkit) v2.4.2 (Mapleson *et al*., 2016) and QUAST v5.0.2 (Gurevich *et al*., 2013). Gene annotation was performed with BRAKER3 v3.0.1 (Gabriel *et al*., 2024), integrating RNA-seq data from the four tissues listed above and protein sequences from species of unknown evolutionary distance. Annotation quality was assessed by estimating RNA-seq coverage on gene models using GeMoMa v1.6.2 (Keilwagen *et al*., 2016; 2018) and by BUSCO v4.0.6 (Manni *et al*., 2021). Repetitive sequences were identified and a repeat library was built using EDTA (Extensive De novo TE Annotator) v1.9.2 (Ou *et al*., 2019) and RepeatModeler v2.0.1 (Flynn *et al*., 2020). The repeat library was used to annotate the genome assembly with RepeatMasker v4.0.9 (Smit *et al*., 1996; https://www.repeatmasker.org/). Collinearity between the genomes of *P. boveana* and *P. edelbergii* (*P.* sect. *Sphondylia*) was assessed using MCScan in the JCVI toolkit (Tang *et al*., 2024). Details are provided in **Methods S2**.

#### Identification, characterization, and expression of Primula boveana S-genes

To assess whether the *S*-locus of the homostylous *P. boveana* differs from those of the heterostylous, closely related *P. edelbergii* (both in sect. *Sphondylia*) and distantly related *P. veris* (sect. *Primula),* we first identified the location and genetic composition of the *S*-locus of *P. boveana* by mapping (i) the nucleotide sequences of *S*-genes from *P. veris* and *P. edelbergii* to the *P. boveana* genome using tBLASTx (--*e*-value 1e-5), and (ii) the gene annotation of *P. edelbergii* onto the *P. boveana* genome using liftoff v1.6.3 (Shumate & Salzberg, 2021). Additionally, we assessed synteny between the *S*-loci of *P. boveana* and *P. edelbergii* from data generated above.

To test whether the *S*-locus of *P. boveana* is present in both haplotypes, as in the *P. edelbergii* genome (Potente *et al*., 2024), we compared the sequencing depth of *P. boveana*’s *S*-locus with the genome-wide average - an approach commonly used to determine whether supergenes occur on one (i.e., hemizygous) or both chromosomes in diploid genomes (Gutiérrez-Valencia *et al*., 2021). Sequencing depth across the *S*-locus and flanking regions was summarized from sliding windows of 10,000 sites from a BAM file with Mosdepth v0.3.1 (Pedersen & Quinlan, 2018). Finally, we compared transposable elements (TEs) between *S*-loci of *P. boveana* and *P. edelbergii* by extracting *S*-locus-specific repeats from the RepeatMasker output (see above).

To determine whether *S*-genes, particularly *CYP^T^*, are disrupted, we compared the gene structures of functional *S*-genes from heterostylous *P. edelbergii* with those of *P. boveana*. Coverage of *S*-genes was estimated for both species using the same approach as for the *S*-locus and flanking regions from sliding windows of 1,000 sites, with the *P. edelbergi*i genome (homozygous at the *S*-locus) as reference. As further validation, coverage of *S*-genes was also estimated from WGS data of a second *P. boveana* individual from a different population (Wadi Shaq Mousa). To identify loss-of function mutations in *S*-genes, *P. boveana S*-gene amino acid sequences were aligned with those of *P. veris* (Potente *et al*., 2022a) and *P. edelbergii* (Potente *et al*., 2024) using MUSCLE v 3.8.31 as implemented in MEGA v11.0.13 (Edgar, 2004; Tamura *et al*., 2021). Using ParaAT v2.0 (Zhang *et al*., 2012), we quantified synonymous and nonsynonymous substitutions and estimated *d_S_*, *d_N_*, and *d_N_*/*d_S_* ratios for each of the four *S*-genes. Finally, the length of *P. boveana*’s *CYP^T^* was assessed by mapping the *P. boveana* assembly to that of *P. edelbergii* using minimap2 v2.24 (Li, 2018; 2021).

To determine whether *S-*genes are expressed, we estimated expression across four tissues (leaf, flower, young floral bud, mature floral bud) from trimmed RNA-seq reads. Transcript expression was estimated with Salmon v1.4.0 (Patro *et al*., 2017), summarized at the gene level and normalized using the median of ratios method (Love *et al*., 2014). Following Hart *et al*. (2013), genes were classified as expressed if normalized RNA-sequence counts were ≥16 (**Fig. S3a**). Details are provided in **Methods S3**.

### Phylogenetic and molecular dating analyses

To resolve the phylogenetic relationship between *P. boveana* within *P.* sect. *Sphondylia*, we generated a chloroplast data set that comprised DNA sequences of four loci (*matK, trnL, trnL– trnF* and *rpl16*) and a total of 17 species: two heterostylous *Primula* species (used as outgroups), nine heterostylous species of *Dionysia*, which is the sister clade of *Primula* sect. *Sphondylia* in the Primulaceae phylogeny of De Vos *et al*. (2014), three heterostylous and three homostylous species of *P.* sect. *Sphondylia*, including *P. boveana*. The only two species of *P.* sect. *Sphondylia* missing from this analysis were the heterostylous *P. davisii* (endemic to a small region in extreme south-east Turkey) and the heterostylous *P. afghanica* (endemic to Afghanistan; Richards, 2003). The chloroplast sequences of all sampled species except *P. boveana* were extracted from the Primulaceae-wide phylogenetic study of De Vos *et al*. (2014) (TreeBASE accession number TB2: S14824). For *P. boveana*, we first assembled its chloroplast genome from WGS data of the reference individual using GetOrganelle v1.7.7.0 (Jin *et al*., 2020) and annotated it with GeSeq (Tillich *et al*., 2017; https://chlorobox.mpimp-golm.mpg.de/geseq.html; accessed April 2026). We then extracted the four loci by extracting *matK, trnL*, and *trnL–trnF* directly from the annotated plastome, and identifying *rpl16 via* BLASTn searches (Camacho *et al*., 2009) with default megablast settings, using the corresponding rpl16 region of *P. edelbergii* (AF402452) as a query. These loci were then incorporated into the Sanger-based alignment of De Vos *et al*. (2014). To identify and mask potential WGS assembly errors, the chloroplast genomes of *P. edelbergii* (SRX24610915) and *P. verticillata* (SRX17744454) were assembled using the same workflow; sites showing discordance between their WGS and Sanger data were masked in the *P. boveana* alignment. Phylogenetic analyses were performed using RAxML-NG v1.2.2 (Kozlov *et al*., 2019) under the GTR+G model with 1000 bootstrap replicates (see Methods S4). We then compared the topology inferred within *P.* sect. *Sphondylia* with the corresponding topology in the dated phylogeny of De Vos *et al*. (2014). Ages of relevant nodes were extracted from De Vos *et al*. (2014), and used them as proxies for nodes of interest in the newly generated phylogeny. Finally, we estimated divergence times between *P. boveana*, *P. edelbergii* and *P. veris* by first estimating synonymous substitution rates (*d_S_*) from genome-wide orthologous genes and subsequently converting these values into absolute divergence ages using a neutral substitution rate of 6.15 × 10⁻⁹ substitutions per synonymous site per year previously determined for *P. veris* (Potente *et al*., 2022a). Details are provided in **Methods S4**.

## Results

### A. Phenotypic analyses

Below, we report results of the analyses aimed at testing whether *P. boveana* is heterostylous or homostylous with developmental variation in herkogamy, and whether this floral morphology represents a delayed-selfing strategy or is rather associated with autonomous selfing conferring reproductive assurance.

#### *Primula boveana* is a homostylous species

sFrequency distributions of anther height and stigma height in 506 flowers sampled after floral organ positions had stabilized, and of pollen size from 19,800 grains from 68 randomly sampled flowers, were unimodal (**Figs 1c**, **S4**), unlike the bimodal distribution expected for heterostylous species. Consistent with this, Hartigan’s dip test detected no significant deviation from unimodality for anther height (*D* = 0.013, *p* = 0.836), stigma height (*D* = 0.016, *p* = 0.546), and pollen size (*D* = 0.002, *p* = 0.999). These results indicate that *P. boveana* is homostylous.

#### Herkogamy changes during anthesis

Herkogamy changed markedly during anthesis in 1,387 flowers sampled between days -2 and +20 (**Figs 1d**, **S2**), as follows: (i) *Initial herkogamy* (days -2 to 0 of anthesis; 159 flowers): 64.8% of flowers exhibited approach herkogamy (stigma above anthers), 29.6% showed no herkogamy (stigma between anthers), and 5.7% had reverse herkogamy (stigma below anthers). (ii) *Herkogamy changes* (days -2 to +6; 842 flowers): Both anther and stigma heights increased during early anthesis (*t* = 13.929, *P* < 0.001; **Fig. 1d**). However, anthers elevated faster than stigmas (*t* = 22.507, *P* < 0.001; **Fig. S5a**; **Table S2a**), resulting in a progressive shift from approach to reverse herkogamy (Fig. 1d). Among 101 plants with ≥3 flowers (798 flowers total), herkogamy negatively correlated with floral age in all cases (Spearman rank test), indicating a consistent developmental pattern. Of these, 56 correlations (55%) were statistically significant at α = 0.05, although correlation strength varied among individuals (**Fig. S6**). (iii) *Final herkogamy* (days +6 to +11; 506 flowers): by day +6, anther and stigma heights had stabilized (**Fig. 1d**). At this stage, 1.4% of flowers exhibited approach herkogamy, 16.8% showed no herkogamy, and 81.8% showed reverse herkogamy. Of the 42 plants that we scored at both flower opening (-2 to 0 days) and maturity (+6 to +11 days), 28 produced flowers spanning more than one herkogamy class at opening, whereas the remaining 14 were uniform for a single class (11 approach, 2 no-herkogamy and 1 reverse herkogamy; **Fig. S7a**). By maturity, all but one plant had converged on reverse herkogamy (**Fig. S7b**). (iv) *Floral withering* (days +12 to +15 and beyond; 149 flowers): Flowers began to wither, and herkogamy fluctuated slightly without a consistent directional trend (**Fig. 1d**).

#### Capacity for autonomous selfing depends on initial herkogamy

Autonomous selfing requires pollen–stigma contact while stigmas are receptive and pollen is available. Pollen germination and tube growth were observed in the stigmas of 15 flowers tested from day 0 to day +6 of anthesis (data not shown). Anther dehiscence began on day -1, increasing from 29.5% of flowers on day 0 to 53.9% on day +1, 83.9% on day +2, 96.9% on day +3, 99.1% on days +4 and +5, and 100% by day +6, when herkogamy stabilized (**Fig. 1d**). These results indicate that stigmas are receptive throughout early anthesis, while pollen availability increases and herkogamy decreases, with both stabilizing on day +6.

Since pollen–stigma contact depends on initial herkogamy and its developmental change, we predicted autonomous selfing to be: (i) highest in flowers with initial approach herkogamy, because anther–stigma contact is achieved after floral opening, when anther dehiscence is near complete or complete; (ii) intermediate in flowers with no initial herkogamy, because anther–stigma contact occurs when anther dehiscence is still low, but anther-stigma distance subsequently increases as dehiscence nears completion; and (iii) lowest in flowers with initial reverse herkogamy, because anther–stigma contact is never achieved (**Fig. S8**). Analyses of autonomous self-pollination in 465 flowers and seed set in 126 flowers (**Fig. S2a**) confirmed these expectations, as follows. In flowers with initial approach herkogamy, 86.6% had pollen on stigmas and 60.0% of ovules developed into seeds. In flowers with no initial herkogamy, 55.0% had pollen on stigmas and 49.1% of ovules developed into seeds. In flowers with initial reverse herkogamy, 23.8% had pollen on stigmas and 24.7% of ovules developed into seeds. All pairwise comparisons among herkogamy classes were significant (*P* ≤ 0.001; **Figs 2**, **S5b**; **Table S2b**). These results showed that the degree of autonomous selfing correlated with initial herkogamy, aligning with our predictions.

**Figure 2.**
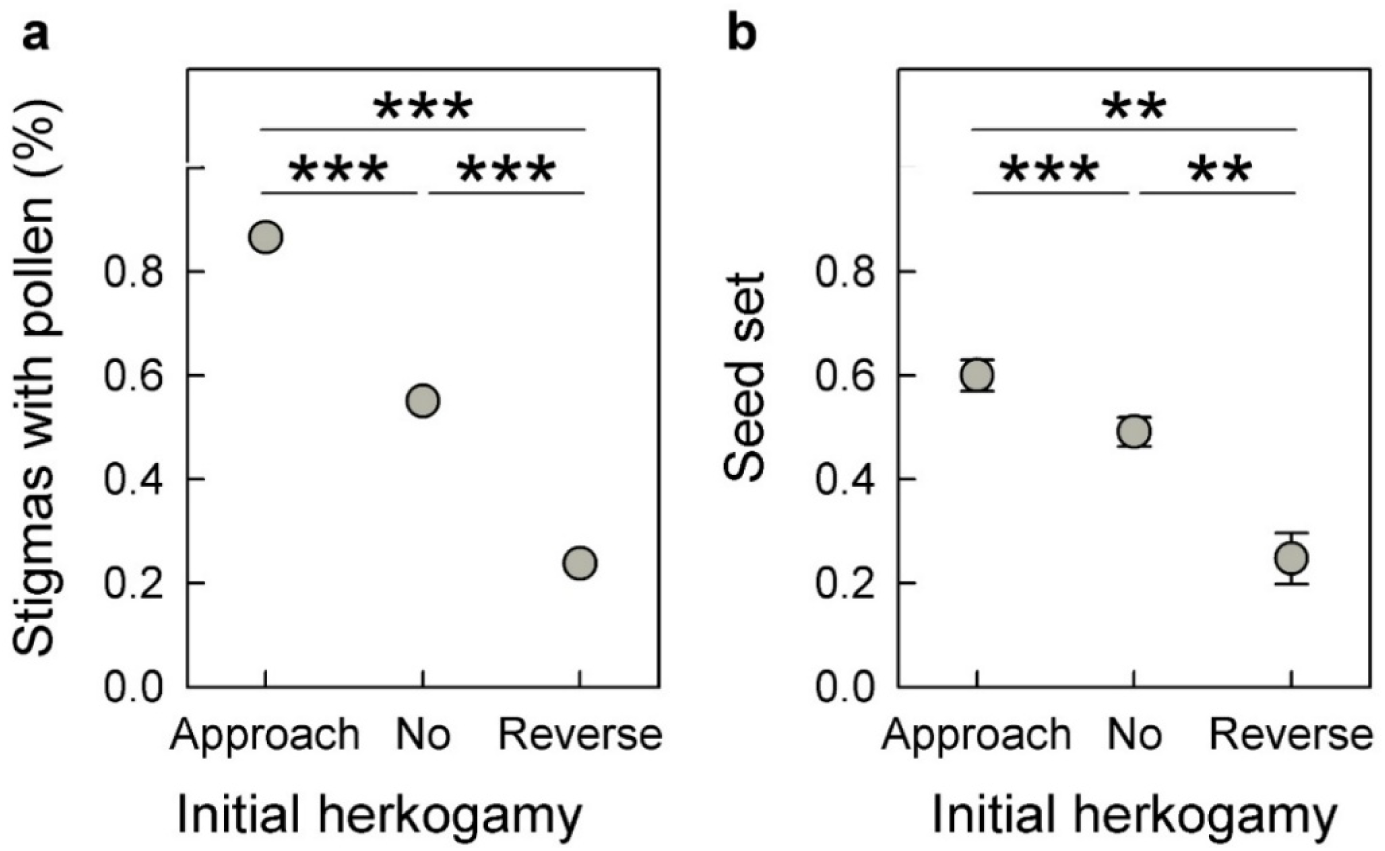
Initial herkogamy and its changes during anthesis determine the timing of anther-stigma contact, thereby mediating selfing opportunities in *Primula boveana*. Comparisons of **(a)** stigmas with pollen (autonomous self-pollination; 465 flowers in total) and **(b)** autonomous seed set (mean values ± standard error; 126 flowers in total) among flowers with approach, no, and reverse initial herkogamy. Sequential Bonferroni correction was used to account for multiple tests. Significance levels: \*\*\**P* ≤ 0.001 and \*\**P* ≤ 0.01. **Panel a:** Sample size per initial herkogamy class: approach = 134, no = 129, reverse = 202; Results of the *χ^2^*test: *χ^2^* = 128.567; df = 2; *p* < 0.001. **Panel b:** Sample size per initial herkogamy class: approach = 47, no = 54, reverse = 25; Results of the generalized linear mixed effects model (GLMM) are provided in Table S2b.

#### Autonomous selfing does not reduce seed set compared to outcrossing

To assess whether autonomous selfing entails a cost to seed production, we compared seed set across three pollination treatments: autonomous selfing, manual selfing, and manual outcrossing. Seed set from autonomous selfing was comparable to, or even exceeded, that from manual selfing and/or outcrossing, despite some population-level variation (**Figs S5c**, **S9**; **Table S2c**). These findings indicate that autonomous selfing can maintain seed production without substantial reduction in quantity.

#### Autonomous selfing provides substantial reproductive assurance

Autonomous seed set differed among flowers with approach-, no-, and reverse-herkogamy (0.600, 0.491, and 0.247, respectively), which corresponded to 81.9%, 67.0%, and 33.7% of the seed set achieved through manual self-pollination (0.733), respectively. When weighted by the frequencies of these classes in the populations (64.8%, 29.6%, and 5.7%), their contributions to overall reproductive assurance were 53.1% (approach-herkogamy), 19.8% (no-herkogamy), and 1.9% (reverse-herkogamy). Overall, autonomous selfing provided 74.8% reproductive assurance, maintaining approximately three-quarters of seed production in the absence of pollinators.

### B. Genomic and phylogenetic analyses

Below we present the *P. boveana* chromosome-scale assembly, the characterization of its *S*-locus, and the phylogenetic analysis that allowed us to place the molecular changes detected at the *S*-locus in a temporal framework.

#### Chromosome-scale genome assembly and comparative genomics

The assembled genome of *P. boveana* was 640.723 Mb (N50 = 66.817 Mb; L50 = 5; **Table 1**), consistent with k-mer-based size estimates (**Fig. S10**). The nine largest scaffolds (64.174-85.355 Mb) contained 99.4% (637.057 Mb) of the assembly, matching the previously reported chromosome number (**Fig. 3**; **Table 1**; Richards, 2003). Assembly quality and completeness were confirmed by k-mer (**Fig. S11**) and BUSCO analyses (92.9% complete genes; **Fig. S12**; **Table 1**). Among the 20,633 identified gene models, 20,168 (97.75%) were supported by RNA evidence (**Fig. S12**; **Table S3**), either partially (2,340 genes; 11.34%) or fully (17,828 genes; 86.41%). BUSCO completeness of the proteome reached 94.9% (**Fig. S12**; **Table 1**). Transposable elements (TEs) accounted for 431.457 Mb (67.3%) of the genome, with LTR retrotransposons (LTR-RTs) comprising 244.858 Mb (38.2%) and being enriched toward chromosome centers (**Figs 3a**, **S13a**; **Table S4**), likely corresponding to pericentromeric regions. Genome-wide heterozygosity was low (0.04%; **Fig. S10**), consistent with elevated inbreeding.

**Table 1:** Statistics for the *Primula boveana* genome assembly and annotation.

|  |  |
| --- | --- |
| Assembly size | 640.72 Mb |
| Number of scaffolds | 226 |
| Cumulative length of the nine largest scaffolds | 637.06 Mb |
| Proportion of the assembly in the nine largest scaffolds | 99.43% |
| N50 | 66.82 Mb |
| L50 | 5 |
| BUSCO complete genes (genome) | 92.9% |
| no. genes | 20633 |
| BUSCO complete genes (proteome) | 94.9% |
| Genome fraction covered by TEs | 67.16% |

**Figure 3.**
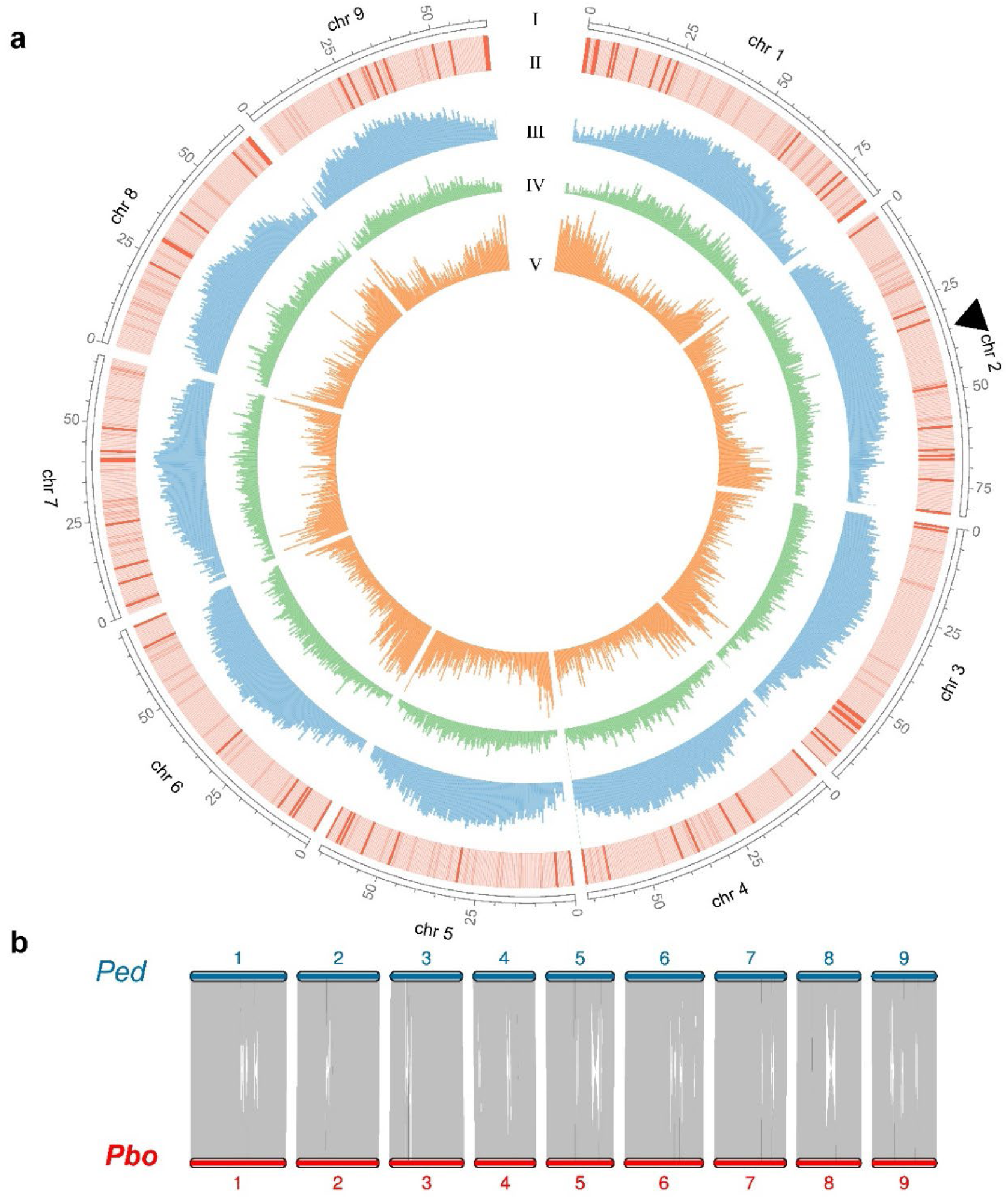
The *Primula boveana* genome and collinearity with *P. edelbergii*. (a) Circos plot of the *P. boveana* genome assembly. Tracks from outside to inside correspond to: (I) the nine chromosomes, with the position of the *S*-locus marked by a black arrow in chromosome 2; (II) GC content, with darker red bars indicating higher GC content; (III) LTR retrotransposons density (blue); (IV) DNA transposons (green); (V) gene density (orange). Tracks II to V were calculated in 500-kb non-overlapping windows. Tracks were visualized using Circos v0.69-9. **(b) Synteny plot of the nine chromosomes of *P. boveana* and *P. edelbergii*** obtained by MCScan. Regions containing >5 collinear genes are connected by grey ribbons.

Comparing the genomes of the homostylous *P. boveana* and the heterostylous *P. edelbergii* revealed strong synteny, with both genomes comprising nine chromosomes and exhibiting similar TE content (67.3% vs. 68.2%) (**Figs 3b**, **S13a**, **S14a**; **Table S4**). The *S*-locus of *P. boveana* shared several key features with that of *P. edelbergii*. In both species, it was located on chromosome 2, contained the same four *S*-genes arranged in identical order (*KFB^T^*, *GLO^T^*, *PUM^T^*, and *CYP^T^*), and had similar TE content (**Figs 3a**, **4a-c**, **S13b**, **Table S4**). Coverage analyses of the reference individual indicated that the *S*-locus is homozygous in *P. boveana* (**Fig. 4a**). In *P. edelbergii*, homozygosity of the reference individual may reflect repeated selfing during cultivation; the *S*-locus was nevertheless inferred by Potente *et al*. (2024) to be hemizygous in wild populations. The following major differences were identified: (i) the *S*-locus of *P. boveana* was inverted and ∼26% smaller than that of *P. edelbergii* (467 kb vs. 633 kb; **Fig. 4b,c**); and (ii) *CYP^T^* was highly degenerated and inverted, being head-to-tail relative to *PUM^T^* in *P. boveana* and tail-to-tail in *P. edelbergii* (**Figs 4d,e**, **S14b**).

**Figure 4.**
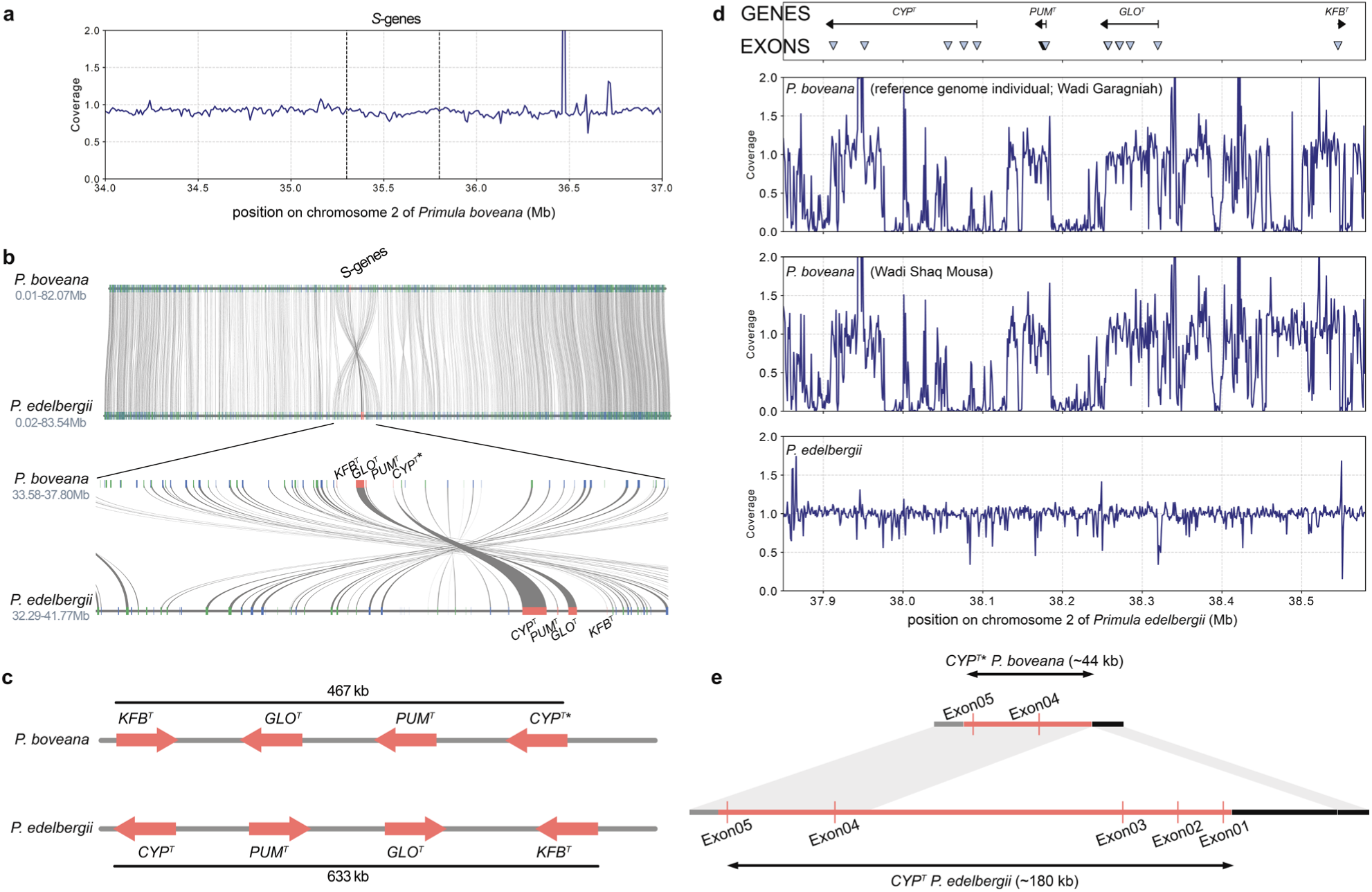
The *Primula boveana S*-locus is homozygous, contains a degraded *CYP^T^*, and is inverted compared to *P. edelbergii*. (a) Sequencing coverage of the genomic region of *P. boveana* chromosome 2 harboring the *S*-genes. (located between the vertical dashed lines; 35.3-35.8 Mb). Coverage normalized by the mean was calculated in 10,000-bp sliding windows. **(b) Microsynteny plot of chromosome 2** of *P. boveana* (top) and its syntenic region in *P. edelbergii* (bottom), with closeup of the region containing the *S*-locus (*S*-genes are shaded in red). (c) Schematic comparison of *S*-locus structure between *P. boveana* and *P. edelbergii. S*-genes are represented as red arrows indicating gene orientation. Gene sizes are not to scale. (d) The *S*-locus of *P. boveana*. Names of *S*-genes and their exon positions and sequencing coverage standardized by the mean (calculated in 1,000-bp sliding windows) of the *S*-locus of *P. boveana* from Wadi Garagniah (top; reference individual) and Wadi Shaq Mousa (middle) and *P. edelbergii* (bottom), as reference. Data for *P. edelbergii* were extracted from Potente *et al*. (2024). (e) Degradation of *CYP^T^* in *P. boveana*. Structure of the disrupted *CYP^T^\** pseudogene in *P. boveana* (top, ∼44 kb, consisting of only 2 exons, as exons 1-3 are missing) and functional *CYP^T^*gene in *P. edelbergii* (bottom, ∼180 kb, consisting of 5 exons). Exon and intron sizes are not to scale. The *P. edelbergii* reference genome was assembled from a short-styled (thrum) individual carrying the dominant *S*-haplotype.

#### *CYP^T^* pseudogenization is consistent with long homostyly

The *P. boveana S*-genes *GLO^T^, KFB^T^*, and *PUM^T^* were not disrupted (i.e. equal exon numbers and lengths as *P. edelbergii*, with no premature stop codons or frameshift mutations; **Fig. S15**, **Table S5**) and expressed in at least one floral developmental stage (**Fig. 5**). Conversely, *CYP*^T^ pseudogenized in *P. boveana*, as exons 1-3 are absent due to a 120-kb deletion, a premature stop codon occurred in exon 4, and a 1-bp frameshift mutation was present in exon 5 (**Figs 4d,e, S14b, S15**). This inference was supported by analyses of a second *P. boveana* individual from a different population, which similarly lacked coverage over *CYP^T^* exons 1-3 (**Fig. 4d**). The pseudogenized *CYP^T^* in *P. boveana* was thus reduced to ∼44 kb, about one-quarter the length of its functional homolog in the heterostylous *P. edelbergii* preventing the identification mutation responsible for the initial loss of heterostyly (**Fig. S14b**). The remaining codon-alignable regions of exons 4 and 5 accumulated 9 synonymous and 9 nonsynonymous substitutions relative to *P. edelbergii*, and 66 synonymous and 42 nonsynonymous substitutions relative to *P. veris* (**Table S5**). Finally, *CYP^T^* was not expressed in any of the three floral developmental stages of *P. boveana*, consistent with the loss of function in this gene (**Fig. 5**).

**Figure 5.**
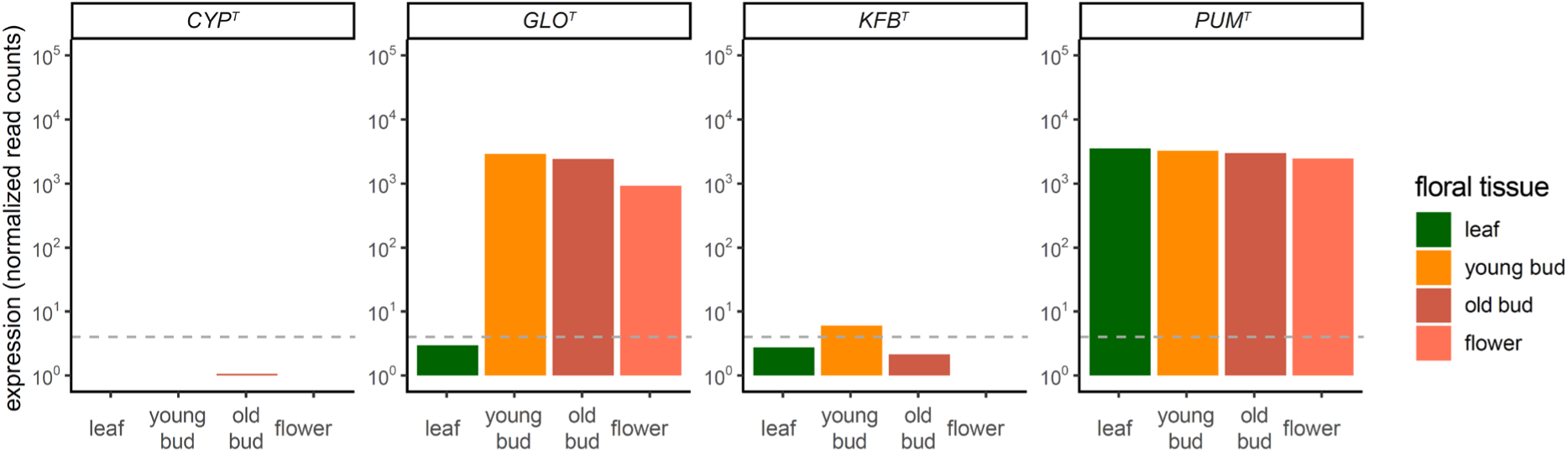
Expression profiles of *S*-genes in *Primula boveana*. Barplots show the number of normalized RNA-seq reads per *S*-gene. Dashed horizontal lines indicate the threshold value at which genes appear not to be expressed (normalized counts > 4; see Fig. S3 and Potente *et al*., 2022b; 2024).

#### Phylogenetic relationships and divergence ages

The maximum-likelihood phylogeny (**Fig. 6b**) strongly supported the monophyly of *Primula* sect. *Sphondylia*, placing *P. boveana* in the same clade as the homostylous *P. simensis* and *P. verticillata*. This homostylous clade was sister to the heterostylous clade formed by *P. gaubeana* and *P. floribunda*, whereas the heterostylous *P. edelbergii* was sister to the rest of *P*. sect. *Sphondylia*. Genome-wide synonymous divergence (*d_S_*) peaked at 0.03 between *P. edelbergii* and *P. boveana* (both *P.* sect. *Sphondylia*) and at 0.24 between *P. veris* (*P.* sect. *Primula*) and *P. boveana* (**Fig. 6a**), corresponding to estimated divergence times of approximately 2.76 Ma (95% CI: 2.57-3.04 Ma) and 19.26 Ma (95% CI: 17.90-21.16 Ma), respectively. The estimated divergence time between *P. edelbergii* and *P. boveana* falls within the confidence interval previously inferred for the split between *P. edelbergii* and the *P. simensis-P. verticillata* lineage (5-95% posterior quantiles: 3.29-9.32 Ma; De Vos *et al*., 2014). Likewise, the estimated divergence between *P. veris* and *P. boveana* falls within the previously inferred confidence interval for the divergence between *P. veris* and the *P. simensis-P. verticillata* lineage (13.86-23.7 Ma) and is consistent with the estimate of 21.95 Ma (± 6.50 SD) reported for the split between *P. edelbergii* and *P. veris* by Potente *et al*. (2024). Together results indicate that the transition to homostyly is shared between *P. boveana*, *P. simensis*, and *P. verticillata* and occurred between 0.93–2.76 Ma.

**Figure 6.**
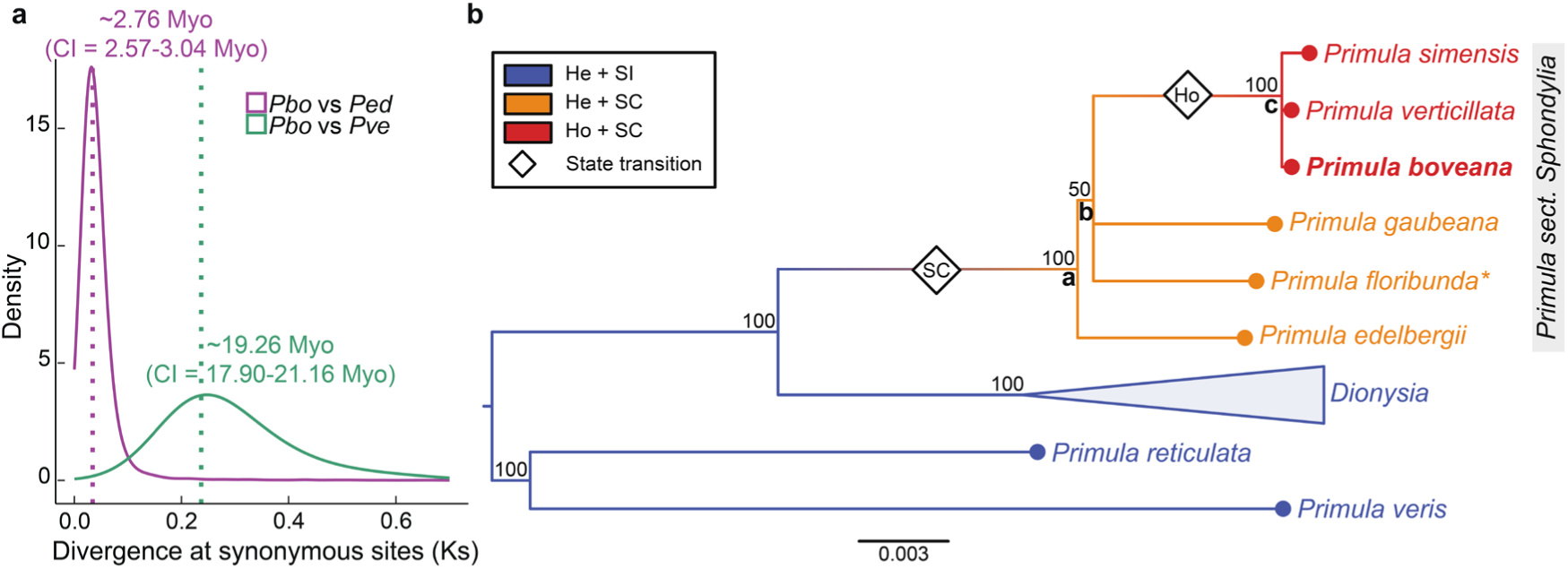
Evolutionary history of *Primula boveana*. **(a) Genome-wide divergence time** estimates based on synonymous substitution rates (*d_S_*). Density distributions of *d_S_* values for collinear orthologues are shown, with dotted lines indicating distribution peaks and corresponding divergence-time estimates (Ma). The estimated divergence between *P. boveana* and the heterostylous *P. edelbergii* (2.57–3.04 Ma) is consistent with the age of node c in (a). **(b) Maximum-likelihood phylogeny of *Primula* sect. *Sphondylia* and its sister clade *Dionysia*** inferred from concatenated plastid loci (*matK*, *trnL*, *trnL–trnF* and *rpl16*). *Primula boveana* forms a clade with the other homostylous species in the section (*P. simensis* and *P. verticillata*). Numbers above branches indicate bootstrap support values ≥ 50%. Divergence times for selected nodes were approximated from (De Vos *et al*., 2014; age Median, 5-95% posterior quantiles): node a = 5.698 Ma (3.291-9.324); node b, 4.259 Ma (2.407-7.356); node c, 0.931 Ma (0.282-2.787). Abbreviations: Ho = homostylous, He = heterostylous, SI = self-incompatible, SC = self-compatible. Asterisk (*) We note that some *P. floribunda* populations are homostylous (Al Wadi & Richards, 1993), but they represent an independent within-species loss of heterostyly.

## Discussion

Our phenotypic and genomic data show that *P. boveana* is homostylous, supporting earlier descriptions (Ernst, 1949; Wendelbo, 1961), but contrasting with later claims of heterostyly (Al Wadi & Richards, 1993). Initial herkogamy and its developmental changes determine the probability of anther-stigma contact in flowers, thereby influencing the probability of autonomous selfing **(Figs 1, 2**). Our results support the hypothesis that long homostyly in *P. boveana* stems from the disruption of the gene controlling style length (**Figs 4, 5**), as observed in other homostylous species within *Primula* (Huu *et al*., 2016; 2022; Mora-Carrera *et al*., 2023; 2024; Mora-Carrera *et al*., 2025) and across angiosperms (Matzke *et al*., 2021; Fawcett *et al*., 2023; Raimondeau *et al*., 2024).

### Developmental changes in herkogamy shape opportunities for outcrossing and selfing

Variation in herkogamy modulates opportunities for outcrossing and selfing in hermaphroditic self-compatible plants (Barrett & Shore, 1987; Belaoussoff & Shore, 1995; Opedal, 2018), with the timing of anther-stigma contact relative to pollen availability, stigma receptivity, and pollinator availability jointly affecting the proportion of selfed and outcrossed seeds (Lloyd, 1992; Lloyd & Schoen, 1992; Schoen & Lloyd, 1992; Blüml *et al*., 2022). In homostyles, flowers with reverse or no herkogamy promote selfing, while those with approach herkogamy may outcross if visited by pollinators (De Vos *et al*., 2012; 2018; Zeng *et al*., 2024). Consistent with the latter, *P. boveana* produces showy flowers typical of insect-pollinated species (Richards, 2003; Omar, 2014a) and exhibits approach herkogamy at anthesis (**Figs 1, S7, S**8). However, herkogamy declines markedly during floral development, resulting in anther-stigma contact in most flowers (**Fig. 1**). In other self-compatible species, similar developmental changes represent a delayed-selfing strategy by maintaining opportunities for outcrossing early in anthesis before autonomous selfing occurs in older flowers (Chen *et al*., 2009; De Vos *et al*., 2012). For example, in the Alpine, homostylous *P. halleri*, anther–stigma contact occurs in only about half of all flowers and typically not until around the fourth day of anthesis, so delayed autonomous selfing is possible only in ∼50% of flowers (De Vos *et al*., 2012; 2018). Conversely, in *P. boveana*, anther–stigma contact occurs within two days of flower opening in ∼90% of flowers, reducing opportunities for outcrossing and promoting early selfing. These findings are consistent with recurrent selfing, as reflected by *P. boveana*’s high capacity for autonomous seed formation (**Fig. 2**), elevated genome-wide homozygosity (**Fig. 10**), and low genetic diversity (Jiménez *et al*., 2013). Early anther-stigma contact is also consistent with theoretical predictions for species experiencing high reproductive uncertainty (Blüml *et al*., 2022), a condition that may characterize *P. boveana* given its dependence on unreliable snow-fed water sourcses and unpredictable pollinator service (Jiménez *et al*., 2013; Zaghloul *et al*., 2016; Abdelaal *et al*., 2020). On the other hand, recurrent selfing is expected to increase the risk of inbreeding depression and reduce genetic diversity, potentially constraining adaptive potential (Baker, 1966; Charlesworth & Charlesworth, 1987; Charlesworth, 2003; Noël *et al*., 2017; Hartfield & Glémin, 2024).

Together, these results indicate that, unlike other species (e.g., *P. halleri*; De Vos *et al*., 2012), the developmental variation in herkogamy observed in *P. boveana* likely does not represent a delayed-selfing strategy, as this species reproduces almost exclusively via autonomous selfing. Such a reproductive strategy may exacerbate vulnerability of *P. boveana* to increasing aridity (Abdelaal *et al*., 2020), with direct consequences to the species long-term survival, although assessment of their long-term evolutionary and demographic consequences is beyond the scope of this study.

### Effects of relaxed selection on pseudogenes

In the ancestrally heterostylous *P. vulgaris*, long-homostylous individuals evolved repeatedly from short-styled plants carrying loss-of-function mutations in *CYP^T^* (Mora-Carrera *et al*., 2023; 2024). In *P. boveana*, homostyly is associated with a 120-kb deletion at the 5’ end of *CYP^T^* that removed exons 1 to 3, a premature stop codon occurred in exon 4, and a 1-bp frameshift mutation was present in exon 5 (**Figs 4, S14b, S15**). Consequently, no evidence of *CYP^T^* expression was detected in all three floral developmental stages of *P. boveana* (**Fig. 5**). Conversely, *GLO^T^*, *PUM^T^*, and *KFB^T^* maintained conserved protein sequence and expression patterns relative to heterostylous *P. edelbergii*. A similar pattern occurs in the homostylous *P. grandis*, where *CYPᵀ* is disrupted (Mora-Carrera *et al*., 2025), supporting the conclusion that *CYP*ᵀ inactivation underlies transitions to long homostyly across recent and older evolutionary contexts. Our results indicate that the transition to homostyly is associated with *CYP^T^*disruption also when such transition occurs in a self-compatible background.

Once a loss-of-function mutation occurs, relaxed selection allows mutations to accumulate (Lahti *et al*., 2009; Cheetham *et al*., 2020) and, over time, non-functional genes may progressively degrade or eventually be lost. In *P. vulgaris*, homostyly evolved 10,000– 12,000 years ago (Mora-Carrera *et al*., 2023; 2024), while our phylogenetic analyses suggest that homostyly in *P.* sect. *Sphondylia* arose once along the stem lineage of the homostylous clade that includes *P. simensis*, *P. verticillata,* and *P. boveana*. Molecular dating by De Vos *et al*. (2014) places this transition between 0.282-7.356 Ma (**Fig. 6b**). Thus, the transition to homostyly preceded diversification of these species and is older than the intraspecific origins of homostyly documented in *P. vulgaris* (Mora-Carrera *et al*., 2023; 2024). The extensive degeneration of *CYP*ᵀ in *P. boveana* (**Figs 4, S14b, S15**) is consistent with a longer history of relaxed selection following loss of heterostyly (Collin & Miglietta, 2008; Lahti *et al*., 2009; Cheetham *et al*., 2020). Accordingly, *CYP*ᵀ degeneration in *P. boveana* exceeds that reported for *P. vulgaris* homostyles and closely resembles the extensive degeneration observed in *P. grandis* (Mora-Carrera *et al*., 2023; 2024; 2025). Given the central role of *CYPᵀ* in determining style length in *Primula* and its extensive degeneration observed in *P. boveana*, re-evolution of heterostyly through restoration of ancestral *CYPᵀ* function appears unlikely. Together, these findings support the hypothesis that prolonged relaxed selection promotes progressive pseudogene degradation and reduces the evolutionary reversibility of trait loss.

## Conclusions

In this study, we demonstrated that *P. boveana* is homostylous, with a developmental decline in herkogamy that promotes early, near-obligate autonomous selfing rather than the delayed-selfing strategy documented in another homostylous *Primula* species, *P. halleri*. This transition is underlain by extensive degeneration of *CYP^T^*, demonstrating that its disruption is related to homostyly even in a self-compatible lineage. The extent of *CYP^T^* degeneration, comparable to that observed in *P. grandis*, together with the old age of homostyly in sect. *Sphondylia*, points to a long history of relaxed selection on this gene that has likely made the back-transition to heterostyly irreversible. Beyond clarifying the genetic basis of homostyly in a self-compatible, diploid background, these findings characterize the reproductive biology of a critically endangered species reliant on autonomous selfing, information directly relevant to its future conservation management.

## Acknowledgements

We thank our research gardeners, Rayko Jonas and Markus Meierhofer, for taking care of the experimental plants and the Functional Genomics Center Zurich (FGCZ) of University of Zurich for providing the infrastructures and support in DNA/RNA sequencing with Illumina platforms. This work was partially supported by the Swiss National Science Foundation (Grant no. 31003A_175556 to EC and P500-3_239422 to GP).

## Authors contributions

B.K. and E.C. conceived the idea and designed the project. H.M. collected the seeds in the natural habitats of *P. boveana*; B.K. performed the pollination experiments and collected and analyzed morphological and reproductive data; B.K. performed the long-read sequencing with the help of N.Y.; B.K. assembled the reference genome with the help of N.Y. and G.P; R.S. provided the short-read data of the additional plant from Wadi Shaq Mousa; B.K., G.P., and E.M.-C. analyzed the genomic data; B.K., E.C., and G.P. wrote the paper with input from all other authors.

## Supporting Information

Additional Supporting Information may be found online in the Supporting Information section at the end of the article.

**Methods S1** Details of the pollination experiments and analysis of stigma receptivity.

**Methods S2** Details of nucleic acid extraction, sequencing, genome assembly and annotation of genes and repetitive elements.

**Methods S3** Details of comparative analysis of the *S*-locus and *S*-gene expression.

**Methods S4** Details of the phylogenetic relationships of species within *P.* sect. *Sphondylia* and divergence times between *Primula boveana*, *P. edelbergii*, or *P. veris*.

**Figure S1** Schematic drawings of floral morphs in heterostylous and homostylous primroses.

**Figure S2** Experimental setup.

**Figure S3** Overview of the *Primula boveana* transcriptome.

**Figure S4** Distributions of anther and stigma heights.

**Figure S5** Evaluation of the performance of the GLMMs.

**Figure S6** Developmental variation in herkogamy per plant.

**Figure S7** Herkogamy within individual plants at flower opening and floral maturity.

**Figure S8** Schematic drawings of developmental changes in herkogamy, anther dehiscence.

**Figure S9** Developmental variation in herkogamy provides reproductive assurance through autonomous selfing.

**Figure S10** Genome size estimates via k-mer analysis.

**Figure S11** Genome assembly quality evaluated by k-mer analysis.

**Figure S12** Genome and proteome evaluation by BUSCO.

**Figure S13** Variation of transposable elements between *Primula boveana* and *P. edelbergii*.

**Figure S14** Synteny between *Primula boveana* and *P. edelbergii*.

**Figure S15** Differences in the protein sequences of the *S*-genes between homostylous and heterostylous species: Amino acid alignments.

**Table S1** Provenance of seeds, seedlings and experimental plants.

**Table S2.** Descriptions and results of GLMMs with contrasts.

**Table S3** GeMoMa results (see FigShare)

**Table S4** Details to transposable elements across the entire genome and the *S*-locus of *Primula boveana*.

**Table S5.** Differences in the protein sequences of the *S*-genes between homostylous and heterostylous species: Synonymous/Non-synonymous substitutions.

