## Supporting Methods, Figures, and Tables for "Phenotypic and genomic changes associated with the evolutionary loss of heterostyly in primroses"

Authors: Authors in alphabetic order, except first and senior author: Barbara Keller, Elena Conti, Hassan Mansour, Emiliano Mora-Carrera, Rebecca L. Stubbs, Narjes Yousefi, and Giacomo Potente

The following Supporting Information is available for this article:

**Methods S1** Details of the pollination experiments and analysis of stigma receptivity.

**Methods S2** Details of nucleic acid extraction, sequencing, genome assembly and annotation of genes and repetitive elements.

**Methods S3** Details of comparative analysis of the *S*-locus and *S*-gene expression.

**Methods S4** Details of the phylogenetic relationships of species within *P.* sect. *Sphondylia* and divergence times between *Primula boveana*, *P. edelbergii*, or *P. veris*.

**Fig. S1** Schematic drawings of floral morphs in heterostylous and homostylous primroses.

**Fig. S2** Experimental setup.

**Fig. S3** Overview of the *Primula boveana* transcriptome.

**Fig. S4** Distributions of anther and stigma heights.

**Fig. S5** Evaluation of the performance of the GLMMs.

**Fig. S6** Developmental variation in herkogamy per plant.

**Fig. S7** Herkogamy within individual plants at flower opening and floral maturity.

**Fig. S8** Schematic drawings of developmental changes in herkogamy, anther dehiscence.

**Fig. S9** Developmental variation in herkogamy provides reproductive assurance through autonomous selfing.

**Fig. S10** Genome size estimates via k-mer analysis.

**Fig. S11** Genome assembly quality evaluated by k-mer analysis.

**Fig. S12** Genome and proteome evaluation by BUSCO.

**Fig. S13** Variation of transposable elements between *Primula boveana* and *P. edelbergii*.

**Fig. S14** Synteny between *Primula boveana* and *P. edelbergii*.

**Fig. S15** Differences in the protein sequences of the *S*-genes between homostylous and heterostylous species: Amino acid alignments

**Table S1** Provenance of seeds, seedlings and experimental plants.

**Table S2** Descriptions and results of GLMMs with contrasts.

**Table S3** GeMoMa results (see FigShare)

**Table S4** Details to transposable elements across the entire genome and the *S*-locus of *Primula boveana*.

**Table S5** Differences in the protein sequences of the *S*-genes between homostylous and heterostylous species: Synonymous/Non-synonymous substitutions

**Methods S1. Pollination experiments and analysis of stigma receptivity**
**Pollination experiments**

Plants were cultivated and pollination experiments conducted in a pollinator-free greenhouse at the University of Zurich. The capacity for autonomous self-fertilization was assessed using intact flowers. For all other treatments, flowers were emasculated before corolla opening by removing the corolla with the attached anthers. Developing fruits were bagged to minimize seed and ovule loss and harvested at maturity. Seeds and undeveloped ovules were recovered from dry capsules and counted from digital images using ImageJ 1.53t. Ovaries contained 279–1161 ovules per flower. Only seven emasculated flowers set fruit, each with few seeds (data not shown), indicating minimal accidental pollination.

**Stigma receptivity test**

The timing of stigma receptivity was assessed by examing pollen grain germination in 15 manually cross-pollinated flowers from seven individuals of the SM population, with floral ages ranging from 0 to 6 days (day 0: 2; day +1: 1; day +4: 4; day +6: 8). Pollen was applied by dabbing dehisced anthers from a different individual of the same population onto the stigma. Flowers were collected 24–48 h after pollination, dissected, and stigmas fixed in 70% ethanol. Pollen tubes were stained and slides prepared following Martin's method (1959), with minor modifications. For each flower, pollen germination was assessed under UV illumination using a fluorescence microscope (BX50, Olympus Corporation, Tokyo, Japan) equipped with a digital imaging system (AxioCam, Axio Vision Rel. 4.8; ZEISS, Oberkochen, Germany). A stigma was scored as receptive if at least one pollen grain had germinated.

**Methods S2**. **Nucleic acid extraction, sequencing, genome assembly and annotation of genes and repetitive elements**
**DNA extraction**

*Wadi Garagniah individual (reference genome individual)* - High molecular weight DNA was extracted from fresh leaves using a modified cetyltrimethylammonium bromide (CTAB) protocol (Healey *et al.*, 2014; Potente *et al.*, 2022) and purified with the Quick-DNA HMW MagBead Kit (Zymo Research). DNA quality and quantity were assessed using Nanodrop (Thermo Fisher Scientific) and Qubit (Invitrogen). Oxford Nanopore libraries were prepared using the LSK-109 ligation kit and sequenced on MinION R9.4.1 flow cells in-house. Basecalling was performed with Guppy v6.6.0.1 using the Super High Accuracy model, yielding 2.63 million reads (43.26 Gb, ~67x coverage; read N50 = 24.21 kb). In addition, Illumina whole-genome sequencing generated 480.01 million paired-end 150-bp reads (72.48 Gb; ~113x coverage; NovaSeq; Functional Genomics Center Zurich, Switzerland). Hi-C data for scaffolding were generated using 219.46 million paired-end 150-bp reads (NovaSeq; Arima Genomics). RNA was extracted from leaves, young floral buds (i.e., petals were closed and did not protrude the calyx), mature floral buds (i.e., petals were still closed but did protrude the calyx), and open flowers (i.e., petals were fully unfolded) using the Spectrum Plant Total RNA Kit (Sigma Aldrich) and sequenced on an Illumina

NovaSeq (~100 million paired-end 150-bp reads per tissue; Functional Genomics Center Zurich, Switzerland).

*Wadi Shaq Mousa individual* - DNA was extracted from silica-dried leaf tissue using the Maxwell Rapid Sample Concentrator 48 and the Maxwell RSC PureFood GMO and Authentication Kit following the manufacturer's instructions without Proteinase K. Whole-genome resequencing was performed on an Illumina NovaSeq, generating ~80 million paired-end 150-bp reads (11.94 Gb; ~18x coverage; Rapid Genomics, Florida, USA).

**Genome assembly**

Genome size was estimated using k-mer frequencies. Forward Illumina reads were used to count 21-mers with the *count* function of Jellyfish v2.2.10 (-C -m 21; Marçais & Kingsford, 2011). The *histo* function of Jellyfish v2.2.10 was used to generate input for GenomeScope (k-mer length = 21; read length = 150; maximum k-mer coverage = 10000; http://qb.cshl.edu/genomescope/; Vurture *et al.*, 2017). Illumina and Nanopore reads were used for de novo assembly with MaSuRCA v4.0.5 (Zimin *et al.*, 2013), with Flye v2.5 (Kolmogorov *et al.*, 2019) used for final contigging. The resulting assembly (641.261 Mb; N50 = 7.016 Mb; L50 = 28) was polished with POLCA (Zimin & Salzberg, 2020) and scaffolded with Hi-C data using YaHS v1.2a.1 (Zhou *et al.*, 2022), improving contiguity (641.302 Mb; N50 = 66.817 Mb;L50 = 4). A Hi-C contact map was generated and visualized in Juicebox (JBAT; Dudchenko *et al.*, 2018) for manual correction, including joining two larger contigs. The final *P. boveana* genome assembly was 640.723 Mb (N50= 66.817 Mb; L50= 5). Assembly quality was evaluated using KAT v2.4.2 (Mapleson *et al.*, 2016), for which the 'histo' file was generated as described above, and BUSCO v4.0.6 (Manni *et al.*, 2021) with the eudicot database (2326 single-copy orthologs; odb v10; www.orthodb.org).

**Gene annotation**

Gene annotation was performed using BRAKER3 v3.0.1 (Gabriel *et al.*, 2024), integrating RNA-seq data from four tissues (leaf, flower, young floral bud, mature floral bud) and protein sequences of species of unknown evolutionary distance. RNA-seq reads were trimmed using Trimmomatic v0.38 (Bolger *et al.*, 2014) with parameters recommended by Trinity (Grabherr *et al.*, 2011): ILLUMINACLIP:2:30:10 SLIDINGWINDOW:4:5 LEADING:5 TRAILING:5 MINLEN:25. Trimmed reads were aligned to the genome assembly (soft-masked using RepeatMasker v4.0.9; https://www.repeatmasker.org/) using Hisat v2.1.0 (Kim *et al.*, 2019) with the parameters --dta --max-intronlen 50'000 --fr. A set of 3'529'642 protein sequences was used, generated by merging the OrthoDB protein data set for Viridiplantae (odb v10; www.orthodb.org) with a high-quality *P. veris* protein set (n=18'900; Potente *et al.*, 2024). BRAKER3 v3.0.1 (Gabriel *et al.*, 2023) was run with default parameters on the *P. boveana* genome assembly using the RNA-seq alignment file and protein dataset as input. Annotation quality was assessed by: (i) assessing the percentage of each gene model supported by RNA-seq was estimated by running ERE and AnnotationEvidence functions of GeMoMa v1.6.2 (Keilwagen *et al.*, 2016; 2018); (ii) using the eudicot database (odb v10), BUSCO v4.0.6 (Manni *et al.*, 2021) was run on the coding sequences of *P. boveana* and *P. edelbergii* (Potente *et al.*, 2024).

**Repeat annotation**
Repetitive sequences were identified and a repeat library constructed using EDTA (Extensive De novo TE Annotator) v1.9.2 (Ou *et al.*, 2019), which combines structure- and homology-based approaches for de novo TE identification. Structure-based TE discovery included LTRharvest (Ellinghaus *et al.*, 2008), LTR_retriever (Ou *et al.*, 2019) for LTRs, TIR-Learner (Su *et al.*, 2019) for TIRs, and Helitronscanner (Xiong *et al.*, 2014) for helitrons. Additional repetitive elements were identified using RepeatModeler v2.0.1 (Flynn *et al.*, 2020). The repeat library was used to annotate the genome assembly using RepeatMasker v4.0.9 (https://www.repeatmasker.org/).

**Methods S3**. **Comparative analysis of the *S*-locus and *S*-gene expression**
**Sequencing depth**

The sequencing depth of the *S*-locus and its flanking regions were estimated from read depth obtained by mapping short reads of the individual used for genome assembly, to the *P. boveana* reference genome. Illumina reads were trimmed using Trimmomatic v0.38 (Bolger *et al.*, 2014; LEADING:3 TRAILING:3 SLIDINGWINDOW:4:15 MINLEN:36) and mapped using BWA-MEM v0.7.17 (Li, 2013). Alignments were sorted, duplicates marked, and indexed using Picard (https://broadinstitute.github.io/picard/) and SAMtools v.1.9–63 (Li, 2011).

**Expression of *S*-genes**

Expression of all four *S*-genes was estimated from four tissues (see above) from trimmed RNA-seq reads. Transcript abundance was quantified using the Salmon v1.4.0 (Patro *et al.*, 2017) *quant* function (mapping-based mode; -gcBias -validateMappings). Salmon output was imported into R v4.3.0 (www. R-project. org/) using Tximport (Soneson *et al.*, 2015; Love *et al.*, 2018), and a A DESeqDataSet was generated with *DESeqDataSetFromTximport* function of DESeq2 (Love *et al.*, 2014) R/Bioconductor package (Gentleman *et al.*, 2004). Expression was summarized at the gene level and gene counts normalized using the median of ratios method (Love *et al.*, 2014).

To classify genes as expressed, distributions of normalized RNA-seq counts were examined across all genes in each tissue. These showed a right skewed distribution, with a higher-expression peak representing actively transcribed genes (Hebenstreit *et al.*, 2011). Following Hart *et al.* (2013), genes were classified as expressed if normalized RNA-seq counts exceeded a conservative threshold of $\geqslant$16 reads, corresponding to the lower bound of this higher-expression peak (Fig. S3).

**Synonymous/Non-synonymous substitution rates**

Pairwise codon alignments and estimates of synonymous substitutions, nonsynonymous substitutions, synonymous and nonsynonymous sites, $d_S$, $d_N$, and $d_N/d_S$ were generated for the four *S*-genes using ParaAT v2.0 (Zhang *et al.*, 2012), which performs sequence alignment using MUSCLE v3.8.31 (Edgar, 2004) and calculates substitution rates using KaKs_Calculator v2.0 (Wang *et al.*, 2010). Because $CYP^{T}$ is pseudogenized in *P.*

*boveana*, only homologous and codon-comparable regions were retained. Codon alignments were generated with ParaAT v2.0 (Zhang *et al.*, 2012) using MAFFT, and $d_N$ (Ka) and $d_S$ (Ks) values were estimated with KaKs_Calculator v2.0 (Wang *et al.*, 2010). A 6 bp region encompassing the premature stop codon at the end of exon 4 and a 30 bp region downstream of the 1 bp frameshift mutation near the end of exon 5 were excluded because these regions could not be reliably compared with the corresponding orthologous regions in *P. edelbergii* and *P. veris*.

**Methods S4**. **Divergence time and phylogenetic relationships between *Primula boveana*, *P. edelbergii*, or *P. veris***

**Phylogenetic relationships**

Plastid sequences of coding (*matK*) and non-coding (*trnL*, *trnL–trnF*, *rpl16*) loci of *P. boveana* were retrieved from plastid sequences assembled from whole genome resequencing (WGS) data of reference genome individual of *P. boveana*, *P. edelbergii* (SRX24610915), and *P. verticillata* (SRX17744454) using GetOrganelle v1.7.7.0 (Jin *et al.*, 2020), which implement a graph-based approach for organelle genome reconstruction. Genome annotation was performed with GeSeq (Tillich *et al.*, 2017; https://chlorobox.mpimp-golm.mpg.de/geseq.html; accessed April 2026), with protein-coding genes, rRNAs, and tRNAs identified using BLAT-based homology searches against the MPI-MP chloroplast reference database. Annotations of protein-coding and rRNA genes were verified using HMMER-based profile searches (Potter *et al.*, 2018) and tRNA genes were additionally predicted with tRNAscan-SE v2.0.7 (Chan *et al.*, 2021) and ARAGORN v1.2.38 (Laslett & Canback, 2004). The *matK*, *trnL*, and *trnL-trnF* regions were extracted directly from the annotated plastomes. The *rpl16* region was identified using BLASTn v2.8.1 (Camacho *et al.*, 2009) with default megablast settings, using the corresponding *rpl16* region of *P. edelbergii* (AF402452) as query. All loci were incorporated into a reduced version of the concatenated alignment of De Vos *et al.* (2014) (TreeBase TB2:S14824), which includes representatives of *Primula* sect. *Sphondylia* (*P. edelbergii, P. gaubeana, P. floribunda, P. simensis*, and *P. verticillata*), nine species of *Dionysia* (*D. hissarica, D. lindbergii, D. aretioides, D. tapetodes, D. revoluta, D. lurorum, D. bryoides, D. haussknechtii,* and *D. gaubae*), and *P. veris* (*Primula* sect. *Primula*) and *P. reticulata* (*Primula* sect. *Aleuritia*) as outgroups. Sequences of *P. boveana* were added to this alignment and manually adjusted in AliView v1.31 (Larsson, 2014) to ensure positional homology. Hypervariable and ambiguously aligned regions were excluded following the curation scheme of De Vos *et al.* (2014). Sites showing discordance between WGS and Sanger data in *P. edelbergii* and/or *P. verticillata* were evaluated in the *P. boveana* alignment, and those present were masked in an alternative dataset. Phylogenetic inference was conducted using RAxML-NG v1.2.2 (Kozlov *et al.*, 2019) under the GTR+G model with 1000 bootstrap replicates. Trees were inferred from both the full and the masked alignments of *P. boveana* to assess the influence of the excluded sites.

**Divergence times**

To estimate divergence times among *P. boveana*, *P. edelbergii* and *P. veris*, we analyzed a separate dataset of collinear nuclear orthologs and estimated synonymous ($d_S$) and nonsynonymous ($d_N$) substitution rates. We identified 18,970, 17,762, and 18,209 orthologous gene pairs between *P. boveana* and *P. edelbergii*, *P. boveana* and *P. veris*, and *P. edelbergii* and *P. veris*, respectively, using MCScan (https://github.com/tanghaibao/jcvi/wiki/MCscan-(Python-version); Tang *et al.*, 2008). Codon alignments and estimates of dN and dS were generated using ParaAT v2.0 and KaKs_Calculator v2.0 as described in Methods S3. Genomic data for *P. edelbergii* and *P. veris* were obtained from Potente *et al.* (2024) and Potente *et al.* (2022), respectively. Divergence times were estimated as $T = d_S/(2r)$, where $r$ is the neutral substitution rate of $6.15 \times 10^{-9}$ substitutions per synonymous site per year (95% CI: $5.60 \times 10^{-9}$ to $6.62 \times 10^{-9}$) previously determined for *P. veris* (Potente *et al.*, 2022).

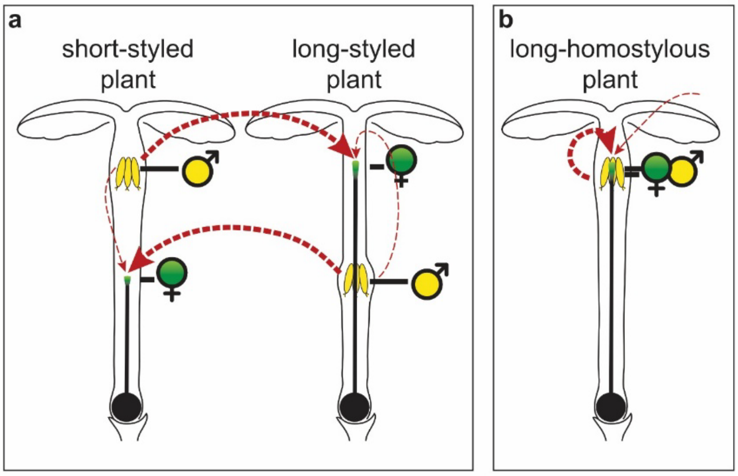


**Fig. S1. Schematic drawings of floral morphs and likelihood of pollen transfer (red arrows) in (a) heterostylous and (b) homostylous primroses. (a)** Heterostylous flowers are either short-styled or long-styled, typically exhibiting exposed and sunken anthers, respectively. Heterostyly optimizes outcrossing by promoting animal-mediated pollen transfer between reciprocally arranged reproductive organs in the two floral morphs, while minimizing pollinator-assisted and diminishing autonomous (i.e., pollinator-unaided) self-pollination between the well-separated anthers and stigmas within the flowers. **(b)** Most homostylous flowers have both anthers and stigmas located at the mouth of the corolla tube (i.e., long homostyly), while short-homostyly is rare (not shown). Spatial separation of anthers and stigmas (i.e., herkogamy) within homostylous flowers is reduced (sometimes zero) and may vary during anthesis (De Vos *et al.*, 2012). The spatial proximity of reproductive organs favors (autonomous) self-pollination, and its timing alters the probability of cross-pollination. The probability of pollinator-assisted pollen transfer between flowers and autonomous pollen transfer is indicated by the thickness of the red arrows. Stigmas are represented in green, anthers in yellow.

(a)

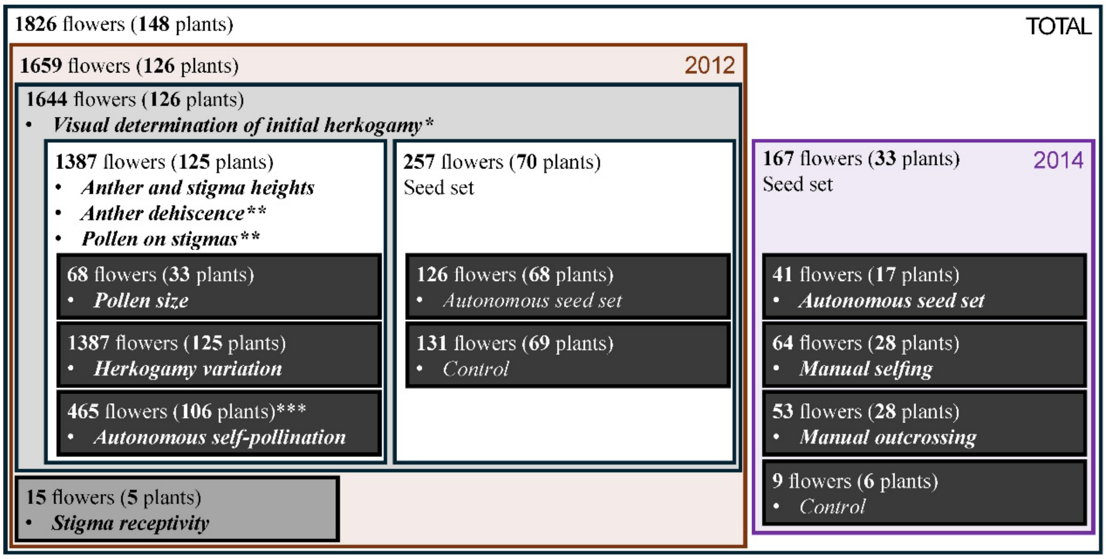


(b)

| | | Day | Approach herkogamy 196 flowers | No herkogamy 373 flowers | Reverse herkogamy 818 flowers | Total 1387 flowers |
|---|---|---|---|---|---|---|
| *Initial herkogamy* (159 flowers) | | -2 | 36 | 7 | 1 | 44 |
| | | -1 | 36 | 12 | 2 | 50 |
| | | 0 | 31 | 28 | 6 | 65 |
| | *Herkogamy changes* (842 flowers) | 1 | 40 | 58 | 20 | 118 |
| | | 2 | 26 | 49 | 42 | 117 |
| | | 3 | 14 | 36 | 52 | 102 |
| | | 4 | 2 | 37 | 74 | 113 |
| | | 5 | 4 | 31 | 88 | 123 |
| *Final herkogamy* (506 flowers) | | 6 | 0 | 14 | 96 | 110 |
| | | 7 | 2 | 16 | 74 | 92 |
| | | 8 | 0 | 12 | 47 | 59 |
| | | 9 | 1 | 20 | 66 | 87 |
| | | 10 | 2 | 11 | 67 | 80 |
| | | 11 | 2 | 12 | 64 | 78 |
| *Floral withering* (149 flowers) | | 12 | 0 | 12 | 42 | 54 |
| | | 13 | 0 | 4 | 30 | 34 |
| | | 14 | 0 | 7 | 21 | 28 |
| | | 15 | 0 | 1 | 7 | 8 |
| | | 16 | 0 | 2 | 4 | 6 |
| | | 17 | 0 | 2 | 9 | 11 |
| | | 18 | 0 | 2 | 2 | 4 |
| | | 19 | 0 | 0 | 2 | 2 |
| | | 20 | 0 | 0 | 2 | 2 |

**Fig. S2. Experimental setup. (a)** Subdivision of experimental flowers into partially overlapping subsets that were used to quantify floral, developmental, and reproductive traits, including autonomous self-pollination and seed set (autonomous, manual self, and manual outcross). Data on anther and stigma heights, anther dehiscence, and stigma receptivity were used to predict outcomes in autonomous self-pollination and seed set, while data on pollen on stigmas was used to estimate autonomous self-pollination. Asterisks: *In 1644 flowers, herkogamy was visually assessed at the time of opening and recorded on a label attached to each flower. As 72 labels became wet during the experiment, the notes were illegible at harvest. Initial herkogamy could therefore be visually assessed in a total of 1572 flowers. **Of 1387 flowers, 92 were excluded, as they had discolored anthers, remained closed, and showed no evidence of pollen release. This left 1295 for analysis. ***From these 1295 flowers, we excluded: (i) 697 harvested 6 days after anthesis (herkogamy still changing); (ii) 108 with stigmas positioned between anthers at harvest (risk of contamination); and (iii) 25 lacking initial herkogamy data. This left 465 for analysis. **(b)** Number of flowers randomly harvested each day during the period spanning two days before and 20 days after floral opening in 2012, subdivided into the three visually determined initial herkogamy categories. Color coding refers to four partially overlapping stages of flower development in *Primula boveana*, defined by flower characteristics that correspond to specific time periods (days): *initial herkogamy* (blue) = the period spanning two days before and the day of flower opening (days –2 to 0 relative to anthesis); *herkogamy changes* (yellow) = the period characterized by developmental changes in herkogamy (days -2 to 6); *stabilized herkogamy* (green) = the period in which positions of anthers and stigmas no longer change, reached on day 6 after flower opening but before any visible signs of withering appear; and *floral withering* (red) = the period in which flowers show visible signs of withering (day 11 onward).

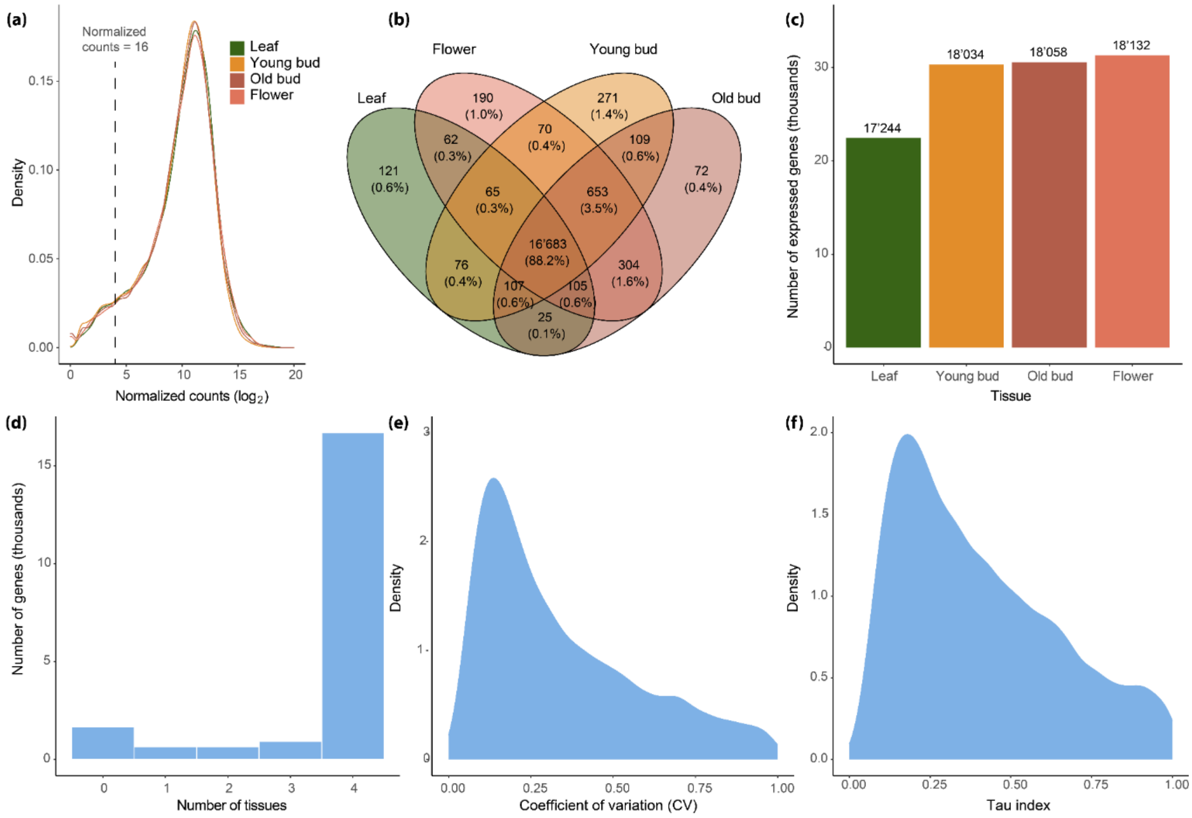


**Fig. S3. Overview of the *Primula boveana* transcriptome. (a)** Density plot showing the distribution of normalized read counts for genes expressed across tissues. The vertical dashed line indicates the threshold above which we marked a gene as expressed in a tissue. **(b)** Venn diagram indicating the number of expressed genes shared by each tissue combination; 16683 genes (88.2%) are expressed in all tissues. **(c)** Number of expressed genes per tissue; floral tissues (reddish hues) are characterized by a higher number of expressed genes than the leaf tissue (green). **(d)** Number of genes expressed in any given number of tissues; most genes are expressed in all four tissues. **(e,f)** Density plots visualize the distribution of two tissue-specificity metrics across all genes, i.e. the coefficient of variation **(e)**, and the Tau index **(f)**.

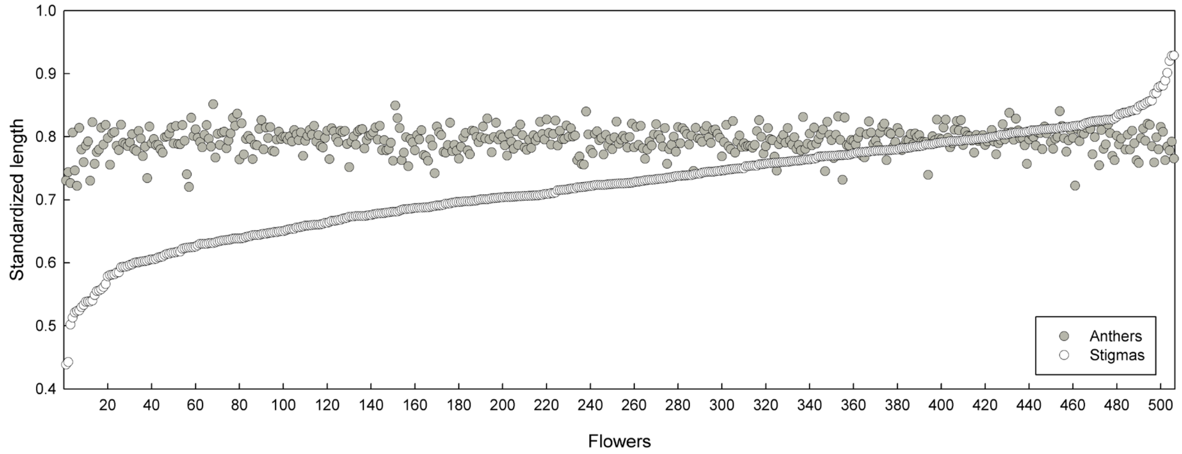


**Fig. S4. Continuous distributions of anther and stigma heights in *Primula boveana*, characteristic of homostylous species.** Anther and stigma heights were measured in 506 mature, non-withered flowers in which positions of anthers and stigmas no longer change (final herkogamy; 6 to 11 days relative to anthesis; see Fig. S1d). Flowers were ranked by stigma height to illustrate the correspondence between the two reproductive organs. Style length and stamen height were standardized by corolla-tube length to control for potential allometric variation in flower size (see Ferrero *et al.*, 2009).

**(a)**

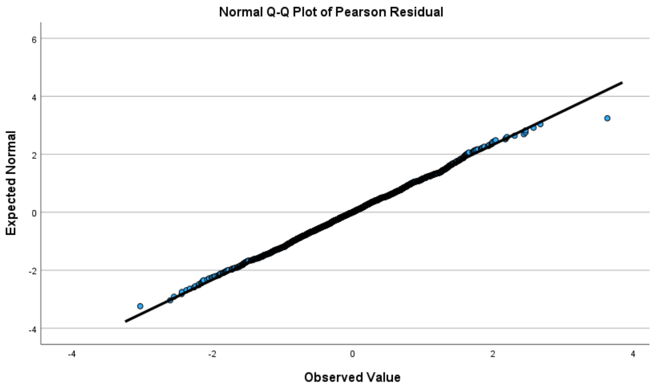


**(b)**

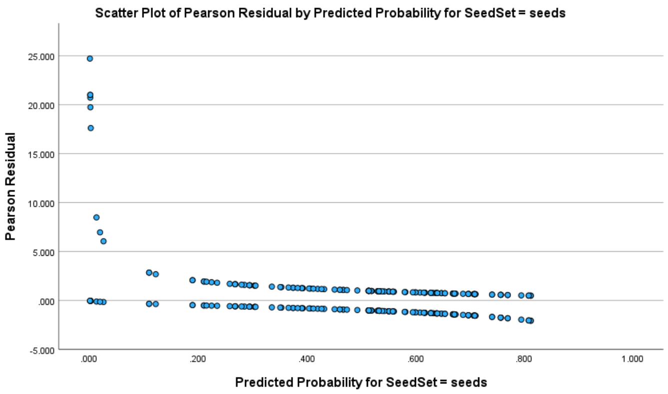


**(c)**

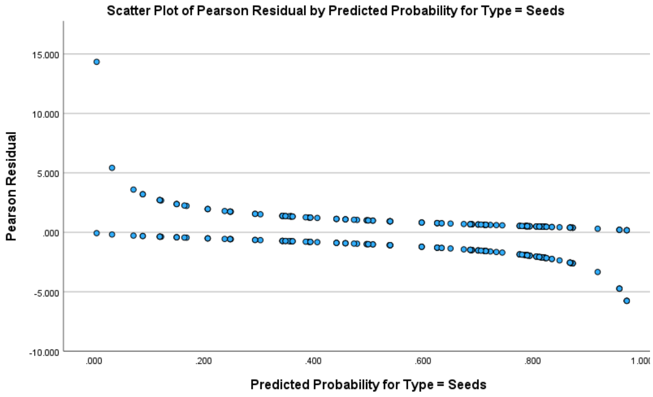


**Fig. S5. Evaluation of the performance of the generalized linear mixed effects models (GLMM)**. **(a)** Normal Q-Q plot assessing normality of Pearson Residuals in the GLMM fitted to test whether anther and stigma positions increased in the first 6 days of anthesis ($\chi^2 = 0.999$; *d.f.* = 1684; $P = 0.543$). **(b, c)** Plots depicting Pearson Residual vs. Predicted Probability, in the GLMM fitted to test whether **(b)** autonomous seed-set differed among initial herkogamy categories and **(c)** autonomous selfing resulted in full seed set.

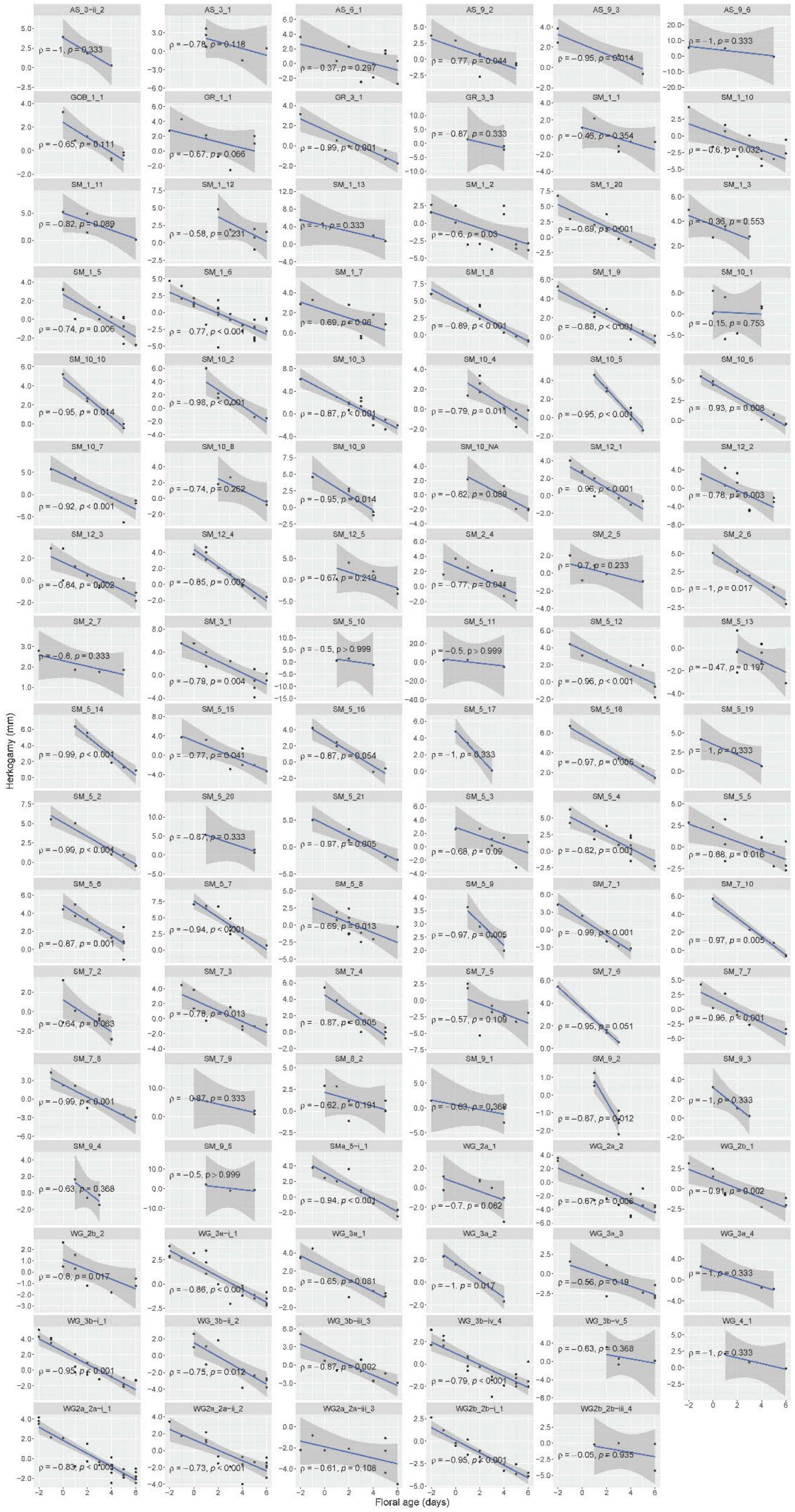

AS_3-ii_2
AS_3_1
AS_6_1
AS_9_2
AS_9_3
AS_9_6
GOB_1_1
GR_1_1
GR_3_1
GR_3_3
SM_1_1
SM_1_10
SM_1_11
SM_1_12
SM_1_13
SM_1_2
SM_1_20
SM_1_3
SM_1_5
SM_1_6
SM_1_7
SM_1_8
SM_1_9
SM_10_1
SM_10_10
SM_10_2
SM_10_3
SM_10_4
SM_10_5
SM_10_6
SM_10_7
SM_10_8
SM_10_9
SM_10_NA
SM_12_1
SM_12_2
SM_12_3
SM_12_4
SM_12_5
SM_2_4
SM_2_5
SM_2_6
SM_2_7
SM_3_1
SM_5_10
SM_5_11
SM_5_12
SM_5_13
SM_5_14
SM_5_15
SM_5_16
SM_5_17
SM_5_18
SM_5_19
SM_5_2
SM_5_20
SM_5_21
SM_5_3
SM_5_4
SM_5_5
SM_5_6
SM_5_7
SM_5_8
SM_5_9
SM_7_1
SM_7_10
SM_7_2
SM_7_3
SM_7_4
SM_7_5
SM_7_6
SM_7_7
SM_7_8
SM_7_9
SM_8_2
SM_9_1
SM_9_2
SM_9_3
SM_9_4
SM_9_5
SMa_5-i_1
WG_2a_1
WG_2a_2
WG_2b_1
WG_2b_2
WG_3a-i_1
WG_3a_1
WG_3a_2
WG_3a_3
WG_3a_4
WG_3b-i_1
WG_3b-ii_2
WG_3b-iii_3
WG_3b-iv_4
WG_3b-v_5
WG_4_1
WG2a_2a-i_1
WG2a_2a-ii_2
WG2a_2a-iii_3
WG2b_2b-i_1
WG2b_2b-iii_4
Herkogamy (mm)
Floral age (days)

**Fig. S6. (previous page). Developmental variation in herkogamy among 798 flowers of 101 plants of *Primula boveana*.** Each scatter plot corresponds to one plant and each circle to one flower. Data shown for plants with ⩾ 3 flowers and floral ages 2 days before to 6 days after flower opening. Blue lines indicate the correlation between herkogamy and floral age per plant (confidence limit indicated = grey shading), with Spearman rho and *p*-value indicated. Note that y-axis ranges vary among individuals.

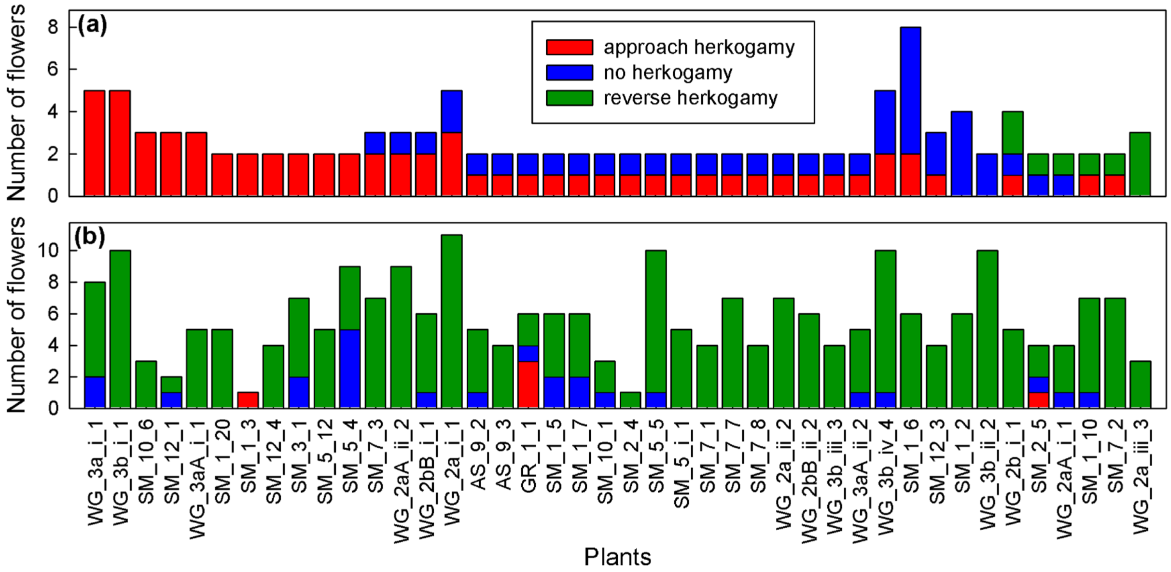


**Figure S7. Herkogamy within individual plants at flower opening (a) and floral maturity (b).** The number of flowers exhibiting approach, no, or reverse herkogamy is shown for 42 plants with ≥2 scored flowers each at flower opening (days −2 to 0) and maturity (days +6 to +11; i.e., flowers whose floral organ positions had stabilized by day 6 of anthesis but are not yet withered; Fig. S2b).

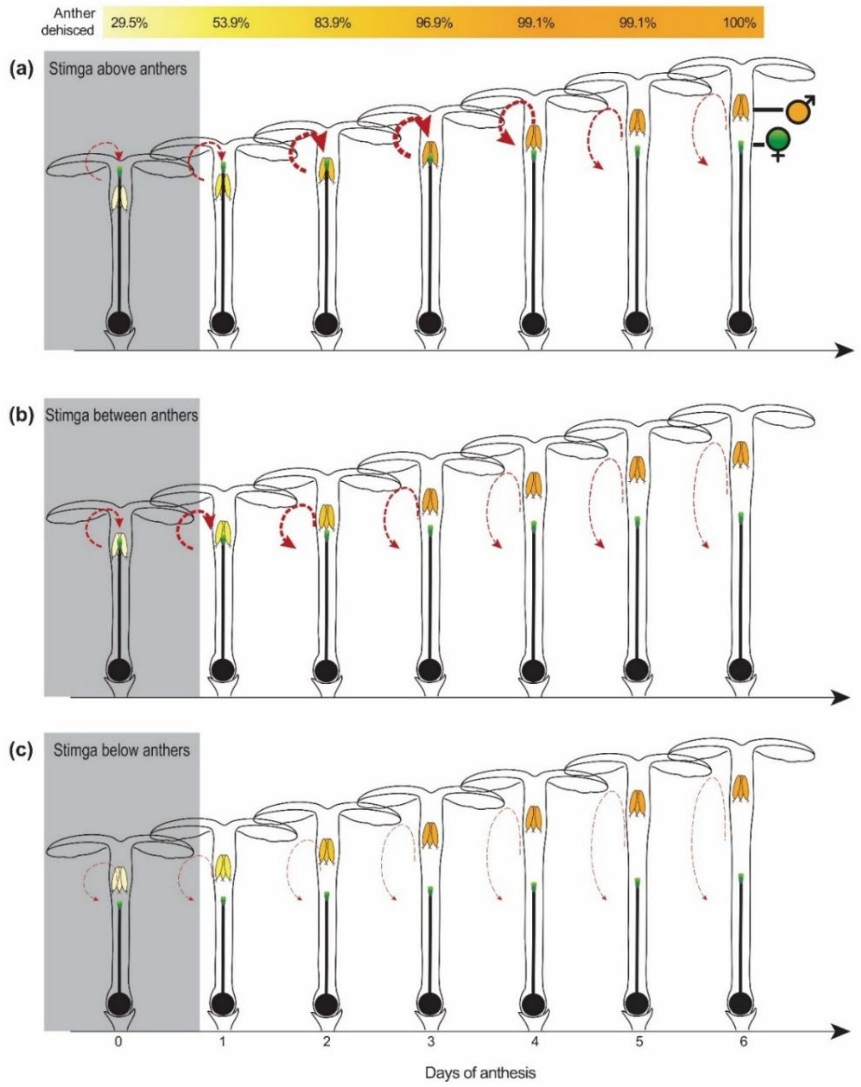


**Fig. S8. Schematic drawings of developmental changes in herkogamy, anther dehiscence (top bar), and likelihood of self-pollination (red arrows).** Among 159 flowers examined around the time of flower opening (floral age between -2 and 0 days; flowers shown in the grey boxes), **(a)** 64.8% had stigmas positioned above the anthers (approach initial herkogamy), **(b)** 29.6% between the anthers (no initial herkogamy), and **(c)** 5.7% below the anthers (reverse initial herkogamy). The probability of self-pollination is indicated by the thickness of the red arrows. Stigmas are represented in green, partially dehisced anthers in light yellow, and fully dehisced anthers in orange. Bar on top indicates the percentage of dehisced anthers (see Fig. 1d). Note anther and stigma positions are not set to scale.

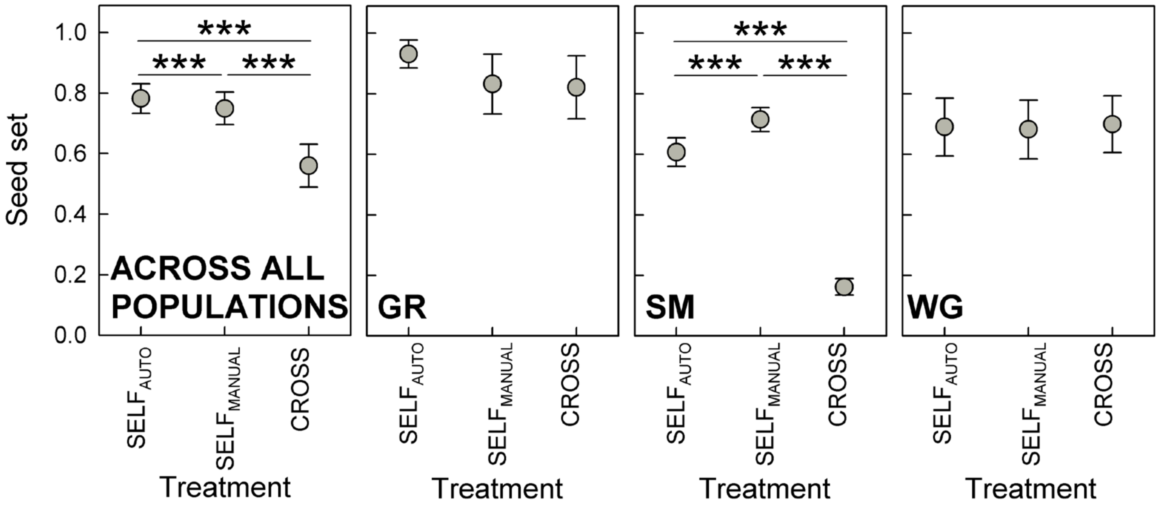


**Fig. S9. Developmental variation in herkogamy provides reproductive assurance through autonomous selfing in *Primula boveana*.** Comparison of seed set (mean values ± standard error) among autonomously selfed ($SELF_{AUTO}$), manually selfed ($SELF_{MANUAL}$), and outcrossed (CROSS) flowers across populations, and within population GR, SM, and WG. Significance levels: ***$P \leqslant 0.001$, all other pairwise comparisons were not significantly different (i.e., $P \geqslant 0.05$). Sequential Bonferroni correction was used to account for multiple tests. Overall, autonomous seed set was significantly higher than seed set in manually selfed and cross-pollinated flowers ($P < 0.001$ for all pairwise comparisons), although there was some variation among populations, as follows: In two populations (GR, WG), autonomous seed set did not differ significantly from either manual selfing or cross-pollination ($P \geqslant 0.223$ for all pairwise comparisons). In the third population (SM), autonomous seed set was high but lower than after manual selfing ($t$ = -11.118, d.f. = 103, $P < 0.001$) and higher than after cross-pollination ($t$ = 20.859, d.f. = 31, $P < 0.001$; Fig. S5c; Table S2c). More detailed studies are needed to elucidate these population-level differences, which fall beyond the scope of the present study.

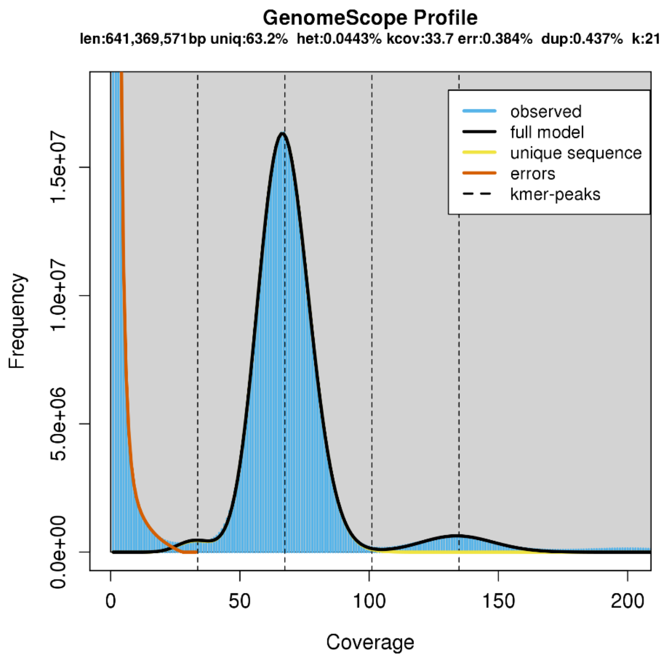


**Fig. S10. Genome size estimates via k-mer analysis.** The genome size estimates for *Primula boveana* using 21-mers corresponded to 641’369’571 bp. Modeling estimated a genome-wide heterozygosity of 0.0443%, corresponding to a homozygosity level of 99.96%, indicating that the *P. boveana* genome is highly homozygous.

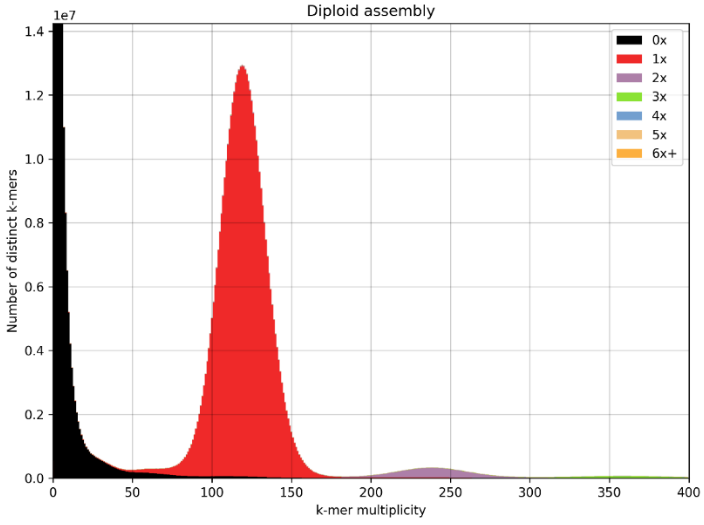


**Fig. S11. Genome assembly quality evaluated by k-mer analysis**. K-mer spectrum (k-mer frequency vs. number of distinct k-mers) obtained with the Kmer Analysis Toolkit (KAT; Mapleson *et al.*, 2016) using Illumina reads . Colors illustrate how many times specific k-mers from the reads appear in the assembly. Red, purple, green, blue, yellow, and orange represent k-mers that appear once, twice, three times, four times, five times, and six or more times in the assembly, respectively. Black represents sequencing errors (i.e. k-mers that are not presented in the assembly). **Red histogram**: The high homozygosity of the genome is reflected by substantially higher peak at k-mer multiplicity of ~125 vs the small heterozygosity peak (at k-mer multiplicity at ~62). **Purple histogram**: A third, short peak at the k-mer multiplicity of ~250 represents genomic regions duplicated in the genome and, as expected, contains k-mers present twice in the assembly.

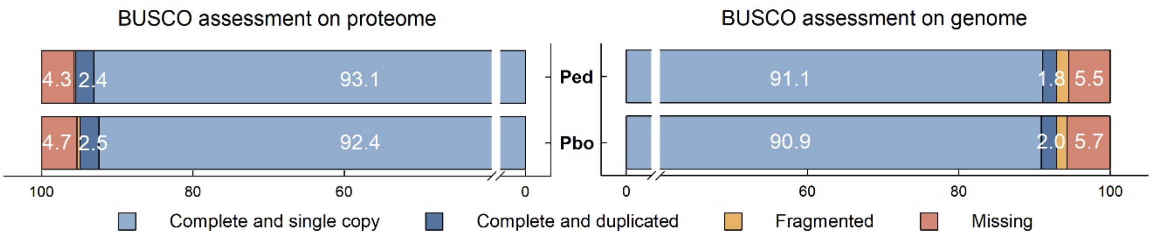


**Fig. S12. Genome and proteome evaluation by BUSCO.** Completeness BUSCO scores for proteomes and genome assemblies of *Primula boveana* (Pbo) and *P. edelbergii* (Ped), obtained using the eudicots_odb10 database, which contains 2326 single-copy orthologs. The total number of complete BUSCO genes for *P. boveana* and *P. edelbergii* is given by the sum of "Complete and single copy" (light blue) and "Complete and duplicated" (dark blue). The percentage for fragmented genes is not reported. Data for *P. edelbergii* was extracted from Potente *et al.* (2024).

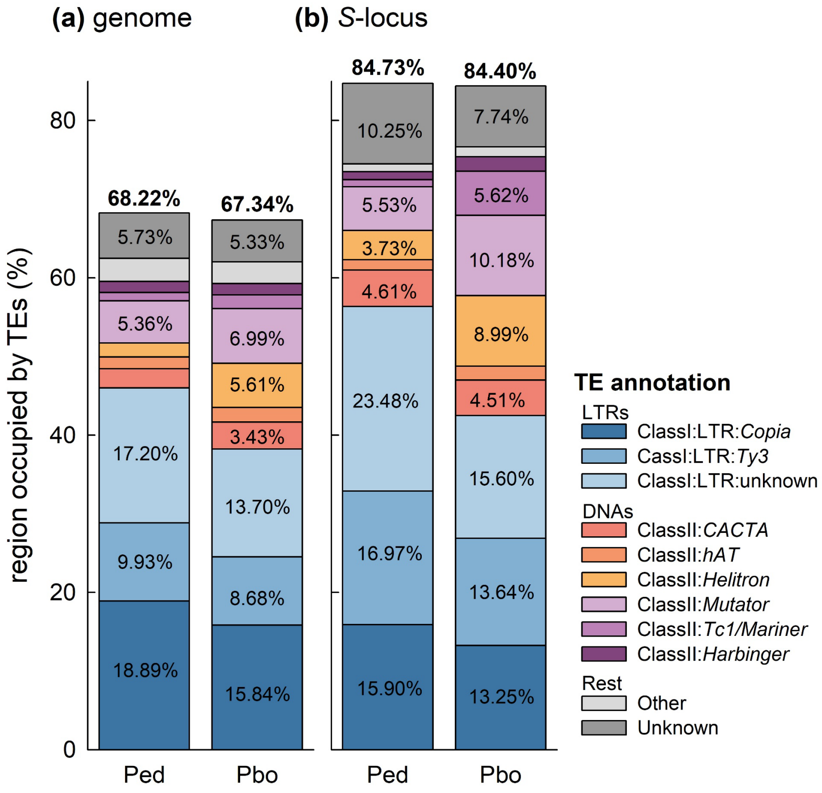


**Fig. S13.** Comparison of **transposable elements (TEs)**. Proportion of TEs in **(a)** the whole genomes and **(b)** the *S*-loci of *Primula boveana* (Pbo) and *P. edelbergii* (Ped). TEs are colored by superfamilies long terminal repeats (LTRs: blue shades) and DNA transposons (DNAs: orange/purple shades). Percentages < 3% are not reported. Data for *P. edelbergii* was extracted from Potente *et al.* (2024).

(a)

(b)

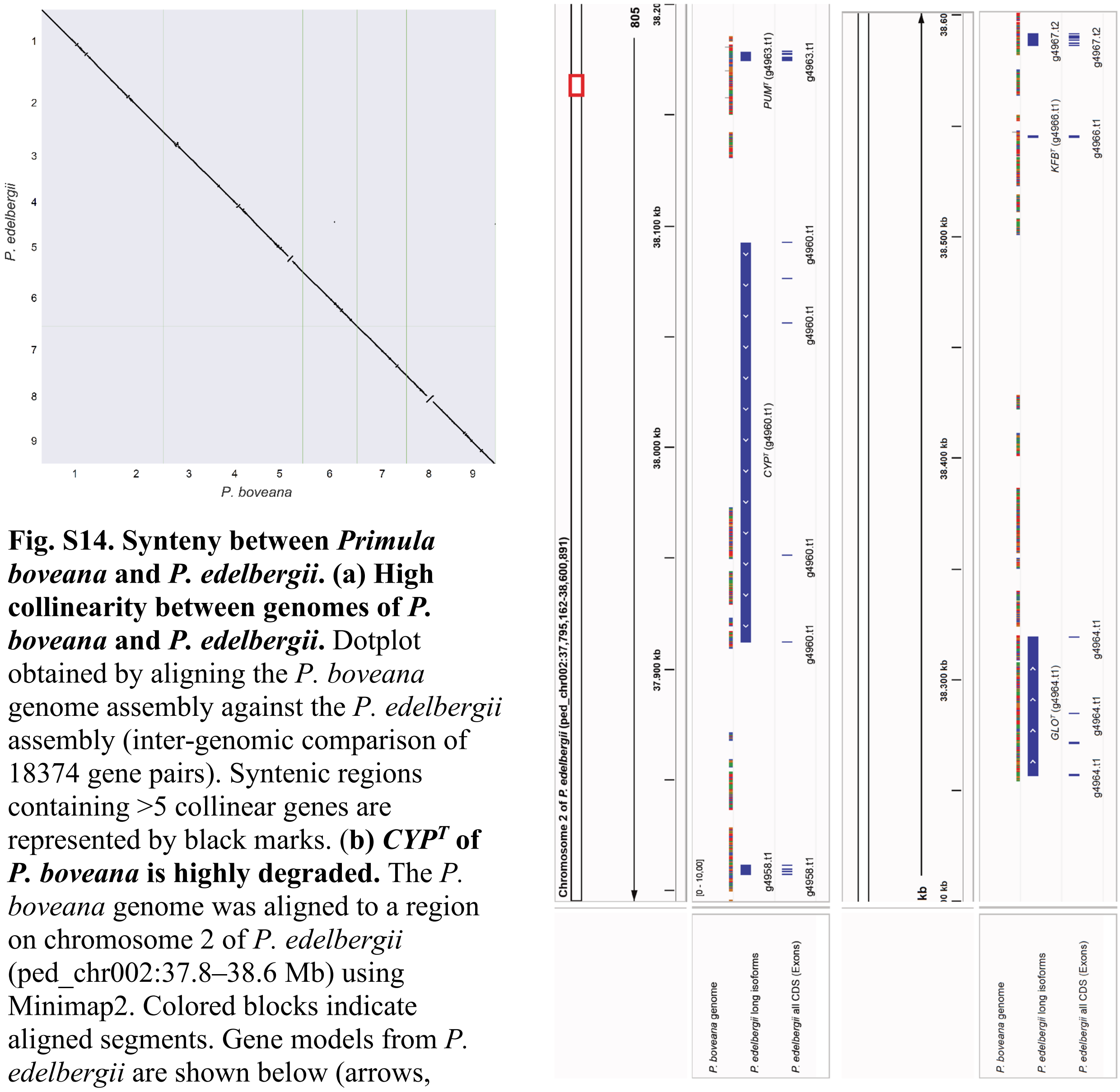


**Fig. S14. Synteny between *Primula boveana* and *P. edelbergii*. (a) High collinearity between genomes of *P. boveana* and *P. edelbergii*.** Dotplot obtained by aligning the *P. boveana* genome assembly against the *P. edelbergii* assembly (inter-genomic comparison of 18374 gene pairs). Syntenic regions containing >5 collinear genes are represented by black marks. **(b) *CYP^T* of *P. boveana* is highly degraded.** The *P. boveana* genome was aligned to a region on chromosome 2 of *P. edelbergii* (ped_chr002:37.8–38.6 Mb) using Minimap2. Colored blocks indicate aligned segments. Gene models from *P. edelbergii* are shown below (arrows, transcripts; bars, exons). Conserved regions show continuous alignments, whereas interruptions indicate divergence or structural variation.

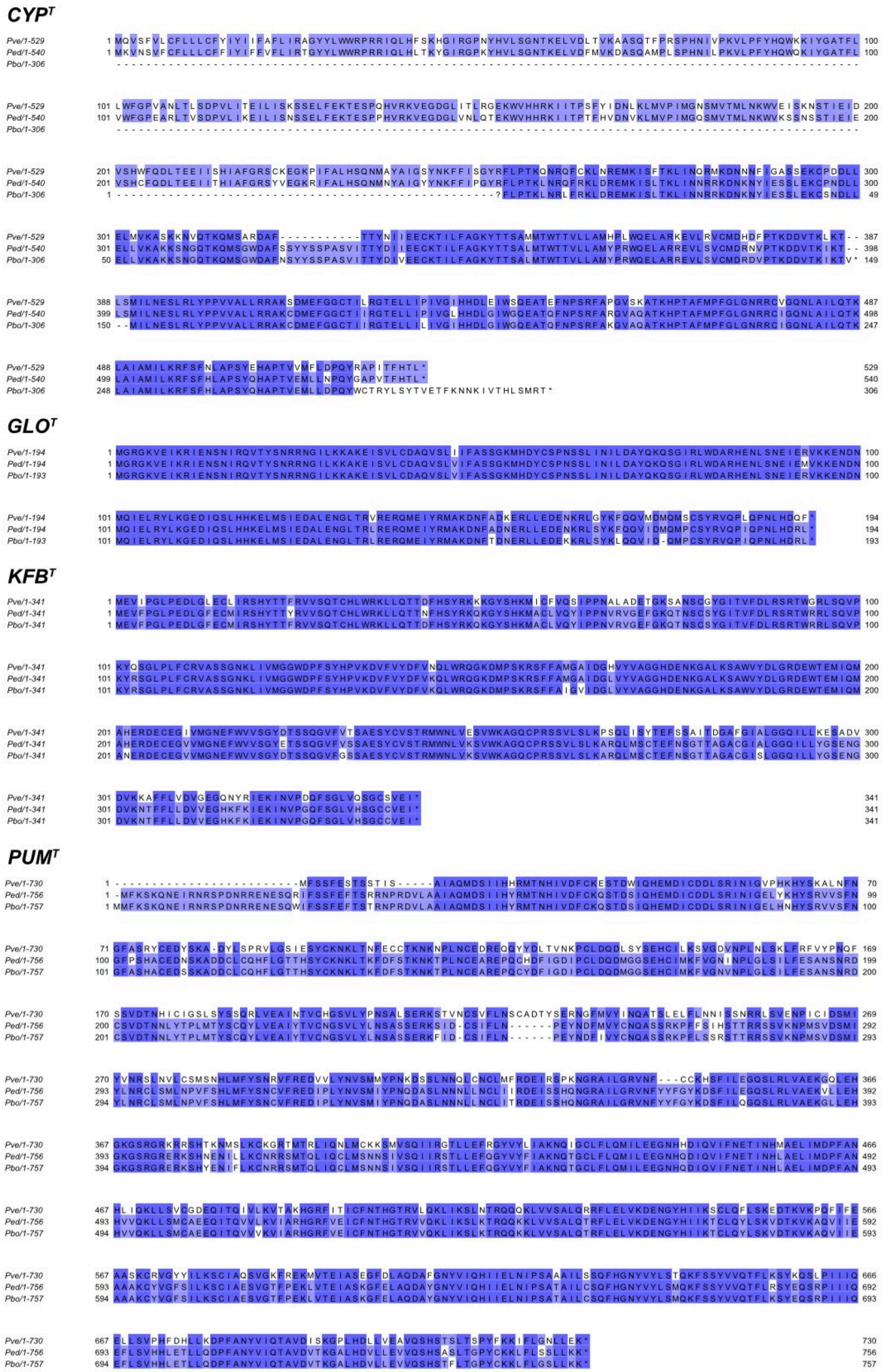
CYP^T
Pve/1-529
Ped/1-540
Pbo/1-306
GLO^T
Pve/1-194
Ped/1-194
Pbo/1-193
KFB^T
Pve/1-341
Ped/1-341
Pbo/1-341
PUM^T
Pve/1-730
Ped/1-756
Pbo/1-757

**Fig. S15. (previous page). Differences in the protein sequences of the *S*-genes between homostylous *P. boveana* (*Pbo*) and those of the heterostylous, closely related *P. edelbergii* (*Ped*) and distantly related *P. veris* (*Pve*) *Primula* sect. *Primula*).** Alignment of the amino acid sequences of the four *S*-genes visualized with Jalview v2.11.3.3 (Waterhouse *et al.*, 2009). Color shading represents percentage identity: Sites with two matches are highlighted in dark purple, sites with one match are highlighted in light purple, and sites with no matches are left white.

**Table S1. Provenance of seeds, seedlings and experimental plants.** Field sampling = Number of natural plants fruits were collected and soil sampled in the field in 2011; Seedlings = number of seedlings transplanted in individual pots; Experimental plants = number of experimental plants used in 2012 and 2014 (see also Fig. S2).

| Population | Field sampling | | | Seedlings | Experimental plants | |
|---|---|---|---|---|---|---|
| | Plants | Soil | Total | | 2012 | 2014 |
| Ain Shennarah (AS) | 12 | 3 | 15 | 64 | 16 | 0 |
| Kahf El-Goulah (GO) | 4 | 1 | 5 | 2 | 1 | 0 |
| Wadi Garagniah (GR) | 4 | 2 | 6 | 13 | 8 | 2 |
| Wadi Shaq Mousa (SM) | 12 | 2 | 14 | 170 | 77 | 26 |
| Wadi Gebal (WG) | 3 | 3 | 6 | 40 | 24 | 5 |
| **Total** | **35** | **11** | **46** | **289** | **126** | **33** |

**Table S2. Descriptions and results of generalized linear mixed effects models (GLMMs) with contrasts**, assessing **(a)** whether anther and stigma positions raise at different rates in the first six days of anthesis, **(b)** whether relative stigma position at beginning of anthesis (above, between, and below anthers) affect capacity of autonomous selfing, and **(c)** whether autonomous selfing results in full seed set across populations, and within populations GR, SM, and WG.

| | Response variable | Error distribution | Link function | Random effects | Source | F | d.f. | *P* | | Contrasts (test statistics, *t*; *d.f.*; adjusted *p*-values) | | |
|---|---|---|---|---|---|---|---|---|---|---|---|---|
| **(a)** | Organ height | Normal | Identity | (i) *field plant* nested within *population*<br>(ii) *experimental plant* nested within *field plant* and *population*<br>(iii) *flower* nested within *experimental plant*, *field plant* and *population* | organ type (O) | 450.628 | 1,840 | < 0.001 | | | | |
| | | | | | days relative to anthesis (D) | 1090.170 | 1,746 | < 0.001 | | | | |
| | | | | | O×D | 506.575 | 1,840 | < 0.001 | | | | |
| **(b)** | Seed set | Binomial | Logit | (i) *field plant* nested within *population*<br>(ii) *experimental plant* nested within *field plant* and *population* | initial herkogamy category | 244.139 | 2, 95347 | < 0.001 | | approach vs. no | approach$_{E}$ vs. reverse | no vs. reverse |
| | | | | | | | | | | 3.996; 39; 0.001 | 3.835; 21; 0.002 | 3.570; 20; 0.002 |
| **(c)** | Seed set | Binomial | Logit | (i) *experimental plant* nested within *field plant* and *population* | treatment (T) | 516.318 | 2,80617 | < 0.001 | | | | |
| | | | | | population (P) | 4.935 | 2,25 | 0.016 | | | | |
| | | | | | T×P | 698.383 | 4,80617 | < 0.001 | | | | |
| | | | | | | | | | | SELF$_{AUTO}$ vs. SELF$_{MANUAL}$ | SELF$_{AUTO}$ vs. CROSS | SELF$_{MANUAL}$ vs. CROSS |
| | | | | | | | | | Across all populations | 4.208; 148; < 0.001 | 9.662; 31; < 0.001 | 10.408; 34; < 0.001 |
| | | | | | | | | | GR | 1.834; 26; 0.223 | 1.859; 26; 0.223 | 0.932; 723; 0.351 |
| | | | | | | | | | SM | -11.118; 103; < 0.001 | 20.859; 31; < 0.001 | 39.061; 32; < 0.001 |
| | | | | | | | | | WG | 0.992; 33678; 0.643 | -0.908; 39553; 0.643 | -1.495; 5892; 0.405 |

$\text{SELF}_{\text{AUTO}}$, autonomous selfing; $\text{SELF}_{\text{MANUAL}}$, manual selfing; CROSS, cross pollination. The models predicted the binary trait seed present/absent correctly in 67.1% and 76.6% of all flowers in **(b)** and **(c)**, respectively. Abbreviations of populations as reported in legend of Fig 1a.

**Table S3.** GeMoMa results (see FigShare).

**Table S4.** Transposable Elements across the entire genome and the *S*-locus of *Primula boveana*.

| | | Genome | | | | *S*-locus | | | |
|---|---|---|---|---|---|---|---|---|---|
| Class / family | | counts | total bp | % of tot TEs | % of genome | counts | total bp | % of tot TEs | % of *S*-locus |
| ClassII:*hAT* | DNA/DTA | 37’864 | 11’930’479 | 2.77 | 1.86 | 27 | 8’259 | 2.1 | 1.8 |
| ClassII:*CACTA* | DNA/DTC | 69446 | 21’955’605 | 5.09 | 3.43 | 57 | 21’060 | 5.3 | 4.5 |
| ClassII:*Harbinger* | DNA/DTH | 38’712 | 9’217’015 | 2.14 | 1.44 | 23 | 8’482 | 2.2 | 1.8 |
| ClassII:*Mutator* | DNA/DTM | 137’021 | 44’760’028 | 10.37 | 6.99 | 129 | 47’509 | 12.1 | 10.2 |
| ClassII:*Tc1/Mariner* | DNA/DTT | 34’363 | 11’102’112 | 2.57 | 1.73 | 36 | 26’230 | 6.7 | 5.6 |
| ClassII:*Helitron* | DNA/Helitron | 87’817 | 35’512’757 | 8.23 | 5.54 | 83 | 41’953 | 10.7 | 9.0 |
| ClassII:*Helitron* | Helitron | 654 | 416’882 | 0.10 | 0.07 | | | 0.0 | |
| Other | LINE/unknown | 2’027 | 1’504’822 | 0.35 | 0.23 | | | 0.0 | |
| ClassI:LTR:*Copia* | LTR/Copia | 121’382 | 101’460’962 | 23.52 | 15.84 | 83 | 61’820 | 15.7 | 13.2 |
| CassI:LTR:*Ty3* | LTR/Gypsy | 60’371 | 55’612’919 | 12.89 | 8.68 | 73 | 63’682 | 16.2 | 13.6 |
| ClassI:LTR:unknown | LTR/unknown | 164’699 | 87’783’687 | 20.35 | 13.70 | 125 | 72’797 | 18.5 | 15.6 |
| Other | Low_complexity | 14’955 | 742’806 | 0.17 | 0.12 | 5 | 276 | 0.1 | 0.1 |
| Other | MITE/DTA | 23’641 | 4’683’487 | 1.09 | 0.73 | 8 | 2’269 | 0.6 | 0.5 |
| Other | MITE/DTC | 4’245 | 543’990 | 0.13 | 0.08 | 1 | 100 | 0.0 | 0.0 |
| Other | MITE/DTH | 7’443 | 1’319’144 | 0.31 | 0.21 | 2 | 185 | 0.0 | 0.0 |
| Other | MITE/DTM | 17’727 | 2’457’795 | 0.57 | 0.38 | 5 | 397 | 0.1 | 0.1 |
| Other | MITE/DTT | 12’439 | 1’860’753 | 0.43 | 0.29 | 5 | 834 | 0.2 | 0.2 |
| Other | Simple_repeat | 79’633 | 3’208’187 | 0.74 | 0.50 | 48 | 1’776 | 0.5 | 0.4 |
| Other | TIR/EnSpm_CACTA | 358 | 276’971 | 0.06 | 0.04 | | | 0.0 | |
| Other | TIR/MuDR_Mutator | 728 | 376’091 | 0.09 | 0.06 | | | 0.0 | |
| Other | TIR/PIF_Harbinger | 725 | 212’552 | 0.05 | 0.03 | | | 0.0 | |
| Other | TIR/Tc1_Mariner | 47 | 12’080 | 0.00 | 0.00 | | | 0.0 | |

| | | | | | | | | | |
|---|---|---|---|---|---|---|---|---|---|
| Other | TIR/hAT | 560 | 379’934 | 0.09 | 0.06 | 1 | 149 | 0.0 | 0.0 |
| Unknown | Unknown | 149’131 | 34’126’438 | 7.91 | 5.33 | 139 | 36’144 | 9.2 | 7.7 |
| Other | pararetrovirus | | | 0.00 | | | | 0.0 | |
| **Total TE** | | **1’065’988** | **431’457’496** | | **67.34** | | **393’922** | | **84.40** |
| Genome/*S*-locus size* | | | 640’723’224 | | | | 466’720 | | |
| Non-Repetitive | | | 209’265’728 | | 32.66 | | 72’798 | | 15.60 |
| total LTRs | | | | 56.75 | 38.22 | | | 50.34 | 42.49 |
| total DNAs | | | | 31.26 | 21.05 | | | 38.97 | 32.89 |
| total MITEs | | | | 2.52 | 1.70 | | | 0.96 | 0.81 |
| total TIRs | | | | 0.29 | 0.20 | | | 0.04 | 0.03 |
| **S*-locus including intronic region before exon04 | | | | | | | | | |

**Table S5. Sequence differences in the *S*-genes between homostylous *Primula boveana* (Pbo), the closely related heterostylous *P. edelbergii* (Ped), and the more distantly related heterostylous *P. veris* (Pve).** Estimates are shown for the numbers of nonsynonymous substitutions (N-subst.), synonymous substitutions (S-subst.), nonsynonymous divergence (*dN*), synonymous divergence (*dS*), and their ratio (*dN/dS*) in pairwise comparisons. **Asterisk:** *Pbo* ($CYP^T$ pseudogene). For comparisons involving *Pbo* in $CYP^T$, analyses were restricted to the remaining portions of exons 4 and 5 that could be aligned. A 6 bp region encompassing the premature stop codon at the end of exon 4 and a 30 bp region downstream of the 1 bp frameshift mutation near the end of exon 5 were excluded because codon homology could not be established unambiguously in these regions. Estimates of *dN*, *dS*, and *dN/dS* were not reported (NR) for comparisons involving *Pbo* because the estimates would be based only on the remaining alignable fragment of a pseudogenized gene and therefore would not be representative of the evolutionary divergence of the intact gene.

| *S*-gene | Comparison | Length (bp) | N-subst. | S-subst. | $d_N$ | $d_S$ | $d_N/d_S$ |
|---|---|---|---|---|---|---|---|
| $CYP^T$ | *Pbo** vs. *Ped* | 828 | 9.18 | 8.82 | NR | NR | NR |
| | *Pbo** vs. *Pve* | 795 | 42.42 | 65.58 | NR | NR | NR |
| | *Ped* vs. *Pve* | 1557 | 96.36 | 99.64 | 0.092 | 0.340 | 0.271 |
| $GLO^T$ | *Pbo* vs. *Ped* | 576 | 3.86 | 5.14 | 0.009 | 0.052 | 0.164 |
| | *Pbo* vs. *Pve* | 576 | 11.66 | 25.34 | 0.031 | 0.223 | 0.140 |
| | *Ped* vs. *Pve* | 579 | 9.43 | 26.57 | 0.026 | 0.227 | 0.114 |
| $KFB^T$ | *Pbo* vs. *Ped* | 1020 | 8.01 | 8.99 | 0.011 | 0.041 | 0.254 |
| | *Pbo* vs. *Pve* | 1020 | 50.00 | 71.00 | 0.078 | 0.343 | 0.229 |
| | *Ped* vs. *Pve* | 1020 | 48.75 | 67.25 | 0.075 | 0.330 | 0.229 |
| $PUM^T$ | *Pbo* vs. *Ped* | 2265 | 23.54 | 11.46 | 0.013 | 0.026 | 0.526 |
| | *Pbo* vs. *Pve* | 2166 | 222.22 | 108.78 | 0.153 | 0.292 | 0.522 |
| | *Ped* vs. *Pve* | 2166 | 222.70 | 103.31 | 0.152 | 0.277 | 0.549 |
